# Aging-Driven Immunosuppression: The Role of Tregs in the Ovarian Tumor Microenvironment

**DOI:** 10.1101/2025.08.07.668717

**Authors:** Mary P Udumula, KM Anjaly, Faraz Rashid, Mohammad Nematullah, Harshit Singh, Tanya Bhardwaj, Miriana Hijaz, Shailendra Giri, Eduardo N Chini E, Heather M Gibson H, Ramandeep Rattan

## Abstract

Epithelial ovarian cancer (EOC) incidence and mortality increase with age, driven in part by chronic inflammation, diminished T cell output, and heightened regulatory T cell (Treg)-mediated immunosuppression. In aged EOC-bearing mice, we observed reduced survival, accompanied by impaired CD4⁺ and CD8⁺T cell responses and a marked expansion of FOXP3⁺ Tregs exhibiting elevated IL-10 and TGFβ expression. Metabolic profiling revealed enhanced oxidative phosphorylation in Tregs from aged mice, along with a fivefold increase in intracellular succinate levels. This accumulation of succinate within the aged tumor microenvironment was found to potentiate Treg suppressive function. Notably, pharmacologic inhibition of α-ketoglutarate dehydrogenase reversed this effect, restoring effector T cell activity. These findings highlight succinate driven metabolic reprogramming as a central mechanism of age related Treg dysfunction in EOC and suggest that targeting succinate metabolism may offer a promising strategy to rejuvenate antitumor immunity in elderly patients.

## Introduction

Epithelial ovarian cancer (EOC), the most lethal gynecologic cancer is a disease of older women, with a median age of 63 years at diagnosis (Langmar & Csomor, 2006). Despite breakthroughs in therapeutic options, EOC’s prognosis remains dismal, with a survival of approximately 50% overall and approximately 20% for patients diagnosed at late stages (stage III/IV) (Torre et al., 2018). Age is a known risk and prognostic factor in EOC, and survival outcomes are worse in older patients than in relatively younger patients, which is further influenced by treatment disparity and severe chemotherapy toxicity (Deng et al., 2017; Tew, 2016; Won et al., 2013). A recent epidemiologic study reported the median survival for women with EOC over the age of 65 years is 37.4 months, whereas it is 47.6 months for women under 65 years (Deng et al., 2017). According to the SEER database, the survival rate for women with EOC is 42% when diagnosed above 65 years while it is 63.7% when diagnosed between 50–65 years, and 73.2% when diagnosed below 50 years of age (National Cancer Institute & Surveillance Epidemiology and End Results Program, 2023). Despite this strong correlation, the molecular pathways that explain aging’s negative impacts on EOC progression are unclear.

Aging is a hallmark of many diseases, including cancer (Anisimov, 2007; Ershler & Longo, 1997) attributed to the predominance of inflammation and associated immune decline (Caruso, Lio, Cavallone, & Franceschi, 2004; Foster, Sivarapatna, & Gress, 2011). With age, the accumulation of DNA damage and genetic mutations resulting from endogenous and exogenous insults leading to genomic instability (Hoeijmakers, 2009), can activate oncogenes or inactivate tumor suppressor genes, thereby promoting malignant transformation (Ou & Schumacher, 2018). Aging reshapes both the innate and adaptive immune systems, resulting in a decline in overall immune competence (Moskalev, Stambler, & Caruso, 2020), contributing to higher susceptibility to cancer. Aging related remodeling of the immune system is called immunosenescence, and is attributed to changes in lymphoid organs, cytokines and a shift in the relative abundance of T-cell subsets (Rodriguez et al., 2020). In this context, aging has recently been associated with increased immunosuppressive regulatory T cells (Tregs; CD4+CD25+FOXP3+), which suppress the number and function of antitumor T cells (Churov, Mamashov, & Novitskaia, 2020; Garg et al., 2014; Jagger, Shimojima, Goronzy, & Weyand, 2014). Tregs suppress immune response via various mechanisms, including production of anti-inflammatory cytokines like IL-10 and TGFβ, by direct cell-to-cell interactions, upregulation of checkpoint proteins (Gonzalez-Navajas et al., 2021; Huang et al., 2015; Huang, Francois, McGray, Miliotto, & Odunsi, 2017; Kamada et al., 2019; Kim, Kim, & Lee, 2020) and metabolic disruption of T cells (C. Li, Jiang, Wei, Xu, & Wang, 2020; Zong, Deng, & Chong, 2024). It plays a key role in EOC (Yigit, Massuger, Figdor, & Torensma, 2010), as their presence in tumor and circulation limits the antitumor activity of effector CD4^+^ and CD8^+^T cells and other immune cells (Cassar, Kartikasari, & Plebanski, 2022; Zhang et al., 2015), resulting in poor prognosis (Kolben et al., 2022; R. Li et al., 2023; L. L. Ye et al., 2020). Previous reports have demonstrated Tregs to increase immunosuppression in aged tumors and play a crucial role in regulating the proliferation of effector T cells (Garg et al., 2014) in melanoma and colon cancer (A. C. Y. Chen et al., 2024; Hurez et al., 2012). This age-related increase in Treg activity may contribute to even greater immune suppression, making immunotherapy less effective in this demographic. Cellular metabolism directs Tregs survival, proliferation, and their suppressive ability (He et al., 2017). In contrast to antitumor effector CD4^+^ and CD8^+^T cells, which primarily rely on glycolysis for their functions (Chang et al., 2013; S. Liu, Liao, Liang, Deng, & Zhou, 2023; Reina-Campos, Scharping, & Goldrath, 2021), Tregs exhibit a distinct preference for oxidative phosphorylation (OXPHOS) (So et al., 2023) for energy, differentiation and expression of anti-inflammatory cytokines, like IL-10 and TGFβ (Chavez & Tse, 2021; Moreno Ayala, Li, & DuPage, 2019). This reliance on OXPHOS allows Tregs to effectively modulate immune responses and maintain immune system homeostasis under normal conditions and in cancer. Thus, understanding the distinct metabolic pathways of the immune cells are crucial for developing targeted therapies aimed at enhancing antitumor immunity. Limited preclinical studies demonstrating the negative impact of age on EOC have revealed underlying mechanisms involving accumulation of senescent cells (Harper, Sheedy, & Stack, 2018), alteration of B cell related pathways in gonadal adipose tissue (Loughran et al., 2018), and ultrastructural changes in collagen of aged omentum (Harper et al., 2022). Our findings validated the rapid and aggressive development of EOC in aged mice as reported by others (Harper et al., 2018; Loughran et al., 2018; Sia et al., 2023). We hypothesize that aging promotes the metabolic reprogramming of Tregs within the ovarian tumor microenvironment, enhancing their immunosuppressive function, impairing effector T cell responses, and thereby accelerating EOC progression in aged mice.

## Materials and Methods

### Cell lines and reagents

The mouse ovarian surface epithelial cancer cell line ID8^p53-/-^ and ID8^BRCA^ ^1-/-^ was a kind gift from Dr. Ian McNeish (Ovarian Cancer Action Research Center, London, UK) (Walton et al., 2016). The mouse ID8 ^p53+/+^, transduced with a lentiviral vector expressing firefly luciferase (ID8-luc2), was kindly provided by Dr. John Liao, University of Washington, Seattle, WA (Liao et al., 2015). The cell lines were maintained in Roswell Park Memorial Institute (RPMI) media (HyClone, Thermo Fisher Scientific, Waltham, MA), supplemented with 10% fetal bovine serum (BioAbChem, Ladson, SC). For in vitro succinate treatments, dimethyl succinate, a cell permeable succinate compound, was purchased from MilliporeSigma (Burlington, VT) (Ehinger et al., 2016), CPI-613 was a generous gift from Cornerstone Pharmaceuticals (Cranbury, NJ). (S)-2-[(2,6-dichlorobenzoyl) amino] succinic acid (AA6) was purchased from MedChem Express (Monmouth Junction, NJ).

### Mice

Female C57BL/6 mice aged 3 months (young) and 22 months (old), representing ∼25 and ∼65 human years (Dutta & Sengupta, 2016), were obtained from Jackson Laboratory (Bar Harbor, ME) and housed under standard conditions at Henry Ford Hospital. All procedures followed institutional animal care and use committee approval (1587). Mice were acclimated for 1 week prior to experiments.

### Tumor induction and monitoring

Mice were injected intraperitoneally (IP) with ID8-luc2, ID8^p53-/-^ and ID8^BRCA^ ^1-/-^ cells (5×10^6^ cells/200 µl Phosphate buffered saline (PBS). Mice were weighed weekly. ID8-luc2 mice underwent bioluminescence imaging weekly as described before using the Xenogen IVIS system 2000 series (Udumula et al., 2021) (Perkin Elmer, Akron, OH). Bioluminescence signals were measured and quantified as total photon flux emission (photons/second) at same exposure time for all mice using Living Image software (Perkin Elmer). For ID8^p53-/-^ and ID8^BRCA^ ^1-/-^ mice, tumor burden was monitored by measuring body weight, abdominal circumference, and ascites formation weekly. Mice exhibiting severe clinical distress, including cachexia, anorexia, labored respiration, excessive ascites accumulation indicated by an abdominal circumference exceeding 8 cm, and those with impaired mobility or compromised bodily functions, were promptly euthanized to ensure ethical compliance and animal welfare (Udumula et al., 2021; Udumula et al., 2023). For CD25 ^+^ T cell depletion, mice with tumor received two IP injections of monoclonal anti-mouse CD25 (IL-2Rα) antibody (100 μg/200 µl PBS; clone PC-61.5.3; BioXcell) (Lebanon, NH) twice weekly (Goschl et al., 2018). Control mice received anti-horseradish peroxidase immunoglobulin G1 isotype control (BioXcell). Depletion was evaluated by flow cytometry quantification of peripheral blood after 6 injections. For survival studies, mice were euthanized when the abdominal circumference reached 8 cm, as per the endpoint criteria approved by the hospital institutional animal care and use committee (protocol-1587). At this stage, mice were humanely euthanized. Mice treated with AA6 received 12.5 mg/kg via IP injection thrice weekly for 4 weeks (Atlante et al., 2018).

### Immune profiling

Tumor single cell suspension was prepared by dissociating tumor cells with Accutase at 37°C for 25 minutes on a shaking platform (Reichard & Asosingh, 2019) (STEMCELL Technologies, Cambridge, MA). The tumor suspension was filtered via a cell strainer and gently washed. The cell pellet was processed for flow cytometry analysis. Lymphocytes from tumors, ascites and blood were also processed for surface marker staining (CD3, CD4, CD8, CD25, CD28, CD57, KLRG1, βgalactosidase and PD1) and intracellular markers (FOXP3, IFNγ, Tbet, granzyme b, Perforin, IL-10, and TGFβ) as before (Udumula et al., 2021; Udumula et al., 2023). Flow cytometric analysis was performed on Attune NxT (Thermo Fisher Scientific), and results were analyzed using FlowJo software (Version 10.9.0; FlowJo, LLC, Ashland, OR). T-Distributed Stochastic Neighbor Embedding clusters were prepared using FlowJo (Version 10.9.0) (Udumula et al., 2023; Zahoor et al.). All fluorochrome antibodies were from BioLegend (San Diego, CA) and listed in Table S1

### T cell suppression assay

Splenic CD4^+^T and CD8^+^T cells were isolated (more than 92% purity) using a MojoSort CD4^+^T cell and CD8^+^T cell isolation kit (BioLegend), labeled with carboxyfluorescein diacetate, succinimidyl ester; 5 mM, as per the manufacturer. Splenic CD4+CD25+FOXP3+ cells were isolated using a MojoSort Isolation Kit (BioLegend) (> 94% purity). CD4^+^T cells and CD8^+^T were co-cultured with Treg cells at a ratio of 2:1 in 200 μl of T cell assay media containing CD3/CD28 Dyna beads and recombinant mouse IL-2 (20 U/mL). After 72-hour flow cytometry was performed for measuring CD4^+^T cell and CD8^+^T cell proliferation (M. L. Chen et al., 2005; Dowling et al., 2018; Geels et al., 2024).

### Quantitation of metabolites by liquid chromatography-tandem mass spectroscopy

Tricarboxylic acid cycle standard mix 1 and 2 (cat no. MSK-TCA1 and MSK-TCA 2 A) and its 13C-labeled metabolites were purchased from Cambridge Isotope Laboratories, Inc. (Tewksbury, MA). Acetonitrile (High Performance Liquid Chromatography grade), formic acid, Mass Spectrometry grade water and methanol were purchased from Sigma-Aldrich (Burlington, VT). The levels of TCA metabolites in various samples were quantified by ultrahigh performance liquid chromatography-tandem mass spectroscopy (Waters, Milford, MA) as described before (Udumula et al., 2024). Glycolysis metabolomic profiling was performed by Metabolon Inc. (Morrisville, MA) (Udumula et al., 2023). Young and old control serum was purchased from George King Bio-Medical (Overland Park, KS).

### Seahorse Metabolic Phenotype

Splenic Tregs from young and aged control and EOC mice were plated (5×10⁵ cells/well) on Cell-Tak coated XFe96 plates, and Oxygen Consumption Rate and Extracellular Acidification Rate were measured using the Agilent Seahorse XFe96 Analyzer as published earlier (Udumula et al., 2024; Udumula et al., 2021; Udumula et al., 2023).

### Single Cell ENergetIc metabolism by profilIng Translation inHibition (SCENITH)

SCENITH was used to quantitatively assess single cell mRNA translation (Arguello et al., 2020). Isolated splenic Treg cells were fixed, permeabilized, and stained with an anti-puromycin antibody followed by treatment with 2-deoxy-D-glucose (2DG, 100 mM), oligomycin (OM, 1 μM), or a sequential combination of these drugs for 45 minutes with puromycin (10 μg/mL) added during the final 30 minutes. Intracellular staining of puromycin was performed using an anti-puromycin monoclonal antibody conjugated to Alexa Fluor 647 (Zahoor et al., 2025) and other surface markers. Quantification of puromycin under various treatments indicates glucose dependence, mitochondrial function, glycolytic capacity, fatty acid oxidation capacity and glutaminolysis (Arguello et al., 2020).

### Western blot

Total protein was isolated, quantified and separated by 10-12% SDS polyacrylamide gel electrophoresis followed by immunoblotting with SUCNR1 antibody (Thermo Fisher Scientific as before (Mangali et al., 2019; Udumula et al., 2022).

### Quantitative-PCR

Total RNA was isolated using the RNeasy kit (Qiagen) (Redwood city, CA), and cDNA was synthesized by iScript cDNA synthesis kit (Bio-Rad) (Hercules, CA) according to the manufacturer’s guidelines. Quantitative PCR analysis was performed using Bio-Rad CFX96 real-time PCR detection system (CFX Maestro, Bio-Rad) as before with ribosomal L27 as housekeeping gene (Udumula et al., 2024; Udumula et al., 2023). Primers sequences are listed in Table S2

### Patient serum studies

Patient blood was collected prior to surgery under an approved institutional review board protocol (# 9520), following informed patient consent and conducted in compliance with the approved methods, guidelines, and regulations. Serum was isolated from the blood of patients under 50 years (young cohort, n = 5) and over 65 years of age (older cohort, n = 5), diluted in complete RPMI media at a 1:25 ratio and then exposed to splenic Treg cells isolated from naive mice. After 48 hours, the Tregs were analyzed for expression of FOXP3, IL-10 and TGFβ by flow cytometry as published before (Udumula et al., 2021; Udumula et al., 2023). For succinate treatments, splenic Treg cell cultures were treated with succinate (100 μM, 1 mM, and 5 mM) for 48 hours followed by flow cytometry analysis.

### 5-Ethynyl-2’-deoxyuridine Staining for proliferation

An EdU Proliferation Assay Kit (iFluor 647, Abcam, Cambridge, UK) (Chapman, Sossick, Boulton, & Jackson, 2012) was used to detect Treg cell proliferation. In brief, 5×10^5^ FITC labelled splenic Tregs isolated from naïve mice were cultured on cover slips in 24 well plates overnight in the presence of young or old ascites along with the culture medium supplemented with 20 mM EdU reagent. After 48 hours, cells were fixed with paraformaldehyde and stained with DAPI. EdU-positive cells were analyzed and quantified by time-lapse fluorescence microscopy (Lonheart FX Automated Microscope, Winooski, VT) and ATTUNE NXT Flow cytometer (Thermo Fisher Scientific).

### ELISA

IL-6, IL-1β, IL-4, IL-10, IFNγ, MCP-1and TNFα levels were measured by ELISA kits from BioLegend. Insulin, IGF-1, Leptin, adiponectin and GMCSF were measured by kits from R&D Systems (Minneapolis, MN), as per manufacturers’ instructions and as published before (Udumula et al., 2023).

### Succinate dehydrogenase assay

Succinate dehydrogenase assay (SDH) activity in splenic Tregs was measured using a colorimetric assay (Abcam) per manufacturer instructions (Almohaimeed et al., 2022).

### Statistical analysis

Statistical tests were performed using GraphPad Prism 8 (Dotmatics, Boston, MA). Data were analyzed by unpaired t-test or one-way analysis of variance as appropriate. Survival curves were evaluated by Kaplan-Meier analysis with Mantel-Cox and Gehan-Breslow-Wilcoxon tests.

Significance was defined as p < 0.05.

## Results

### Aging exacerbates preclinical EOC models

To understand aging’s impact on EOC, we compared the tumor progression and survival of IP injected mouse ID8 EOC cells with different gene signatures including ID8^p53+/+^-luc2, ID8^p53−/−^ and ID8^p53-/-, BRCA^ ^1-/-^ in young and old C57BL/6 female mice (Roby et al., 2000; Walton et al., 2016). Old mice bearing ID8-luc2 tumors had shorter median survival (49 vs. 65 days in young mice) (Figure 1A,J) and showed greater tumor burden, confirmed by increased abdominal circumference, ascites (Figure 1B,C), and higher bioluminescence imaging signals (Figure S1A,B). Older mice consistently weighed more than younger mice throughout the study (Figure S1C,D). The mice with ID8^p53−/−^ and ID8^p53-/-, BRCA^ ^1-/-^ tumors representing high grade serous ovarian cancer (HGSOC) (Mandilaras et al., 2019) showed similar patterns. Old mice with ID8^p53−/−^ tumors had a median survival of 39 days compared to 48 days in young mice (Figure 1D,J), along with increased abdominal circumference and ascites accumulation (Figure 1E,F). Old mice with ID8^p53-/-, BRCA^ ^1-/-^ had a median survival of 33 days compared to 45 days in young mice (Figure 1G,J), and an increased abdominal circumference and ascites volume (Figure 1H,I). These observations are in accordance with other reports (Hou et al., 2022; Loughran et al., 2018; Shim et al., 2022) that demonstrate similar aggravated EOC progression in older mice, thus, confirming aging exacerbates EOC progression. Aging is also associated with increased levels of cellular senescence and inflammatory factors (Bottazzi, Riboli, & Mantovani, 2018; X. Li et al., 2023), defined as senescence-associated secretory phenotype (SASP) that support a tumor permissive microenvironment (Dong et al., 2024; Kuilman et al., 2008; Narita & Lowe, 2005). In accordance, we observed metabolic growth factors IGF-1 (Figure S2B,F) and leptin (Figure S2D,H) levels were significantly increased in the ascites of aged mice compared to young mice bearing ID8^p53-/-^ and ID8 ^p53-/-, BRCA1-/-^ tumors, while insulin (Figure S2A,E) was lower in the aged ID8^p53-/-^ mice compared to young mice but was unchanged in ID8^p53-/-, BRCA1-/-^ mice.

**Figure 1:**
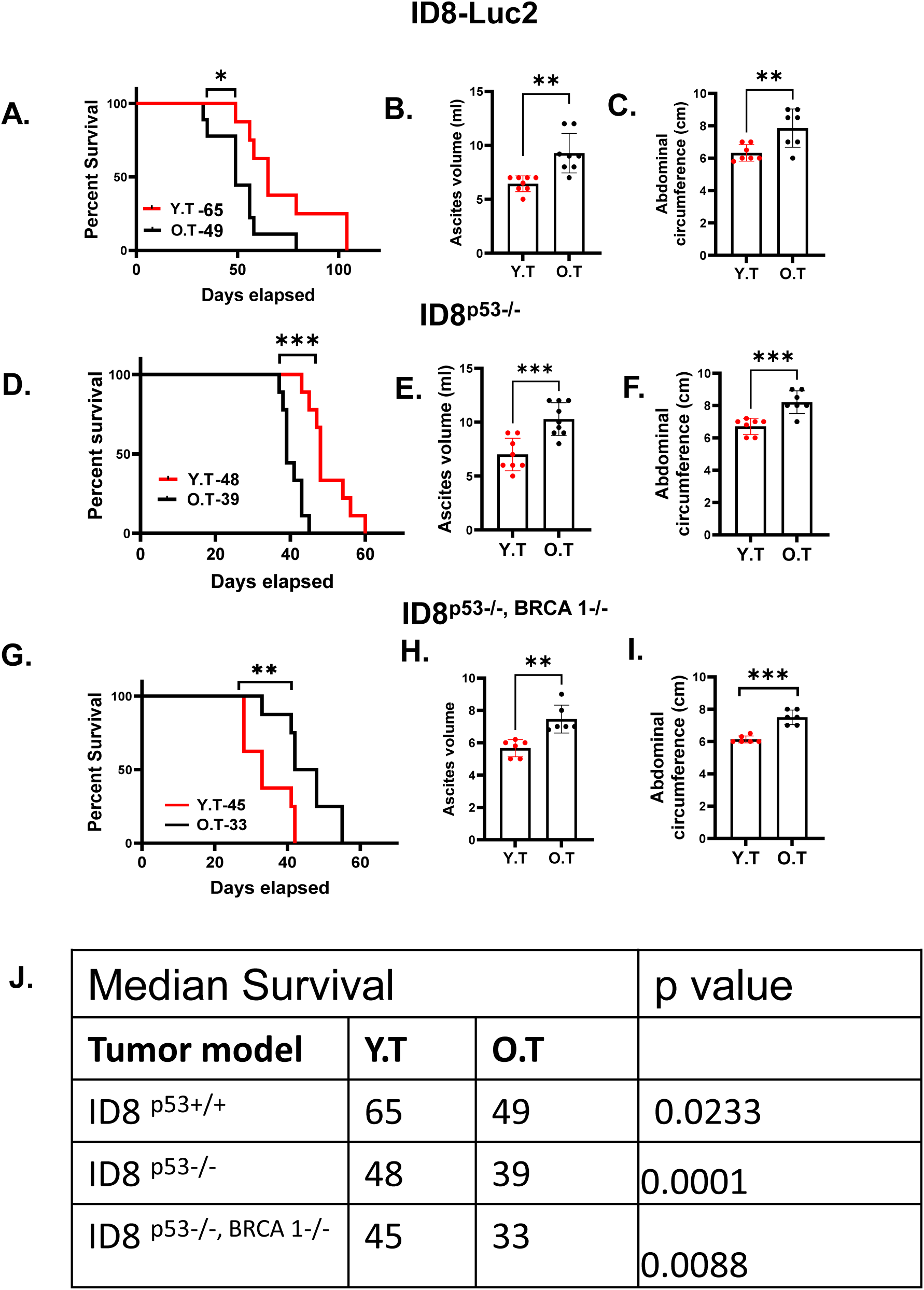
Aging exacerbates in preclinical EOC models: Kaplan–Meier graphs indicating overall survival in mice bearing (A–I) (A) ID8-luc2 EOC injected in Young Tumor (Y.T) or Old Tumor (O.T) (n = 10/group) mice, p = 0.0233 by Gehan-Breslow-Wilcoxon test; (D) ID8^p53−/−^ EOC injected in Y.T or O.T (n = 10/group), p < 0.0001 by Gehan-Breslow-Wilcox test and (G) ID8^p53−/−, BRCA^ ^1-/-^ EOC injected in Y.T or O.T (n = 10/group), p = 0.0088 by Gehan-Breslow-Wilcox test. (B, E, and H) Ascites volume collected at 6 weeks from ID8-luc2, and at 5 weeks from ID8^p53−/−^ and ID8^p53−/−, BRCA^ ^1-/-^ EOC bearing young and old mice. (C, F, I) Bar graph represents average abdominal circumference at 6 weeks of ID8-luc2, and at 5 weeks of ID8^p53−/−^ and ID8^p53−/−, BRCA^ ^1-/-^ EOC bearing young and old mice. (J) The table represents median survival in all three mouse models of young and old EOC mice. *p<0.5, ∗∗p < 0.01, ∗∗∗p < 0.001, OT compared with YT group by Student’s *t* test.

Adiponectin levels were unchanged in the ascites of ID8^p53-/-^ mice but reduced in aged ID8 ^p53-/-, BRCA1-/-^ mice relative to younger mice (Figure S2C,G). Among SASP markers, levels of IL-4 (Figure S2I,M), IL-10 (Figure S2K,O), MCP-1 (Figure S2J,N), IL-1β (Figure S2Q,U), IL-6 (Figure S2S,W), and TNFα (Figure S2R,V) were significantly increased in ascites of aged ID8^p53-/-^ and ID8 ^p53-/-, BRCA1-/-^ mice, compared to young mice. In contrast, GMCSF levels were decreased in both aged models (Figure S2L,P). IFNγ, a key cytokine for cytotoxic T cells, was lower in the ascites of aged mice of ID8^p53-/-^ (Figure S2T) while it was increased in ID8 ^p53-/-, BRCA^ ^1-/-^ mice compared to young EOC mice (Figure S2X). These findings are consistent with other cancer studies indicating that the aging Tumor microenvironment (TME) induces distinct modifications that contribute to the establishment of a pro-tumorigenic milieu (Fane & Weeraratna, 2020; Hibino et al., 2021).

### Aging induces differential systemic T cell response in EOC

Age-related immune decline, particularly in T cell composition, is well-established (Salam et al., 2013; Yager et al., 2008). To examine this, we profiled major immune populations in the blood of young tumor-bearing mice, old tumor-bearing mice, young, and old control mice without tumors. In control old mice without tumors, we found no significant changes in the number of CD4^+^T cells, while the CD8^+^T cell number was significantly decreased and a notably higher prevalence of Tregs was observed in old control mice compared to young controls (Figure S3C,F,E), consistent with previous studies (Deng et al., 2017; Garrido-Rodriguez et al., 2021; M. Li et al., 2019; Palatella, Guillaume, Linterman, & Huehn, 2022; Yang et al., 2024). In EOC-bearing mice, the young mice exhibited an increase in the number of both CD4^+^ and CD8^+^T cells compared to young control mice without tumors (Figure S3A-C,F), along with a robust increase in effector CD4+IFNγ+ (Figure S3D) and CD8+IFNγ+ cells (Figure S3G). Additionally, cytotoxic markers, granzyme B (Grz b) and perforin, were significantly increased in young mice with tumors, highlighting an effective T cell-mediated response against the tumor (Figure S3H,I). In contrast, old mice with EOC showed a marked decline in both CD4^+^ and CD8^+^T cells compared to old controls without tumors (Figure S3A-C,F). The failure to mount a comparable antitumor immune response by old EOC mice was reflected in decreased effector CD4+IFNγ+ and CD8+IFNγ+ cells, and cytotoxic CD8+Grz b+ and CD8+perforin+ cells compared to young EOC mice (Figure S3D,G,H,I). While Tregs were increased in both young and old EOC mice, it was most pronounced in old EOC mice (Figure S3A,B,E), suggesting a heightened Treg-mediated immunosuppression. Analysis of control mice revealed that the percentage of total macrophages (CD11b⁺F4/80⁺) was lower in old mice compared to young counterparts (Figure S3J).

Additionally, there were no significant changes in the expression of M1 (CD38) (Figure S3K) or M2 markers (EGR2 and CD206) between young and old control mice (Figure S3L,M). The co-expression of CD38/EGR2 and CD38/CD206 also remained comparable between the groups (Figure S3N,O). Notably, the M1 macrophage marker iNOS was significantly reduced in aged control mice relative to young controls (Figure S3P). Intracellular M2 markers (EGR2⁺Arg1⁺ and CD206⁺Arg1⁺), as well as myeloid-derived suppressor cells (CD11b⁺Gr1⁺), did not differ significantly between young and aged control mice (Figure S3Q-S). No significant increase in the percentage of F4/80⁺ cells was observed between young EOC-bearing mice and young control mice. In contrast, a significant increase in F4/80⁺ cells were detected in old EOC-bearing mice compared to aged control mice (Figure S3J). The percentage of F4/80+CD38+ cells did not differ significantly between young and old EOC mice compared to their respective young and old control groups (Figure S3K). The percentage of F4/80+EGR2+ cells remained unchanged between young EOC mice and young control mice; however, a significant increase in F4/80+EGR2+ cells was observed in old EOC mice compared to both young EOC and old control mice (Figure S3L). Both young and old EOC mice exhibited an increased frequency of CD206+ and CD11b+Gr1+ populations compared to their age-matched control mice; however, no significant differences were observed between the young and old EOC groups (Figure S3M,S). Additionally, a decrease in the M1/M2 ratio (CD38/EGR2) was observed in older EOC mice compared to young EOC mice, although this ratio did not differ significantly when compared to other groups (Figure S3N). The overall M1/M2 ratio (CD38/CD206) was reduced in EOC mice of both age groups relative to their non-tumor-bearing counterparts (Figure S3O), indicating an increased presence of M2-like macrophages associated with tumor progression.

Additionally, the M1 macrophage marker iNOS showed a significant decline in both young and old EOC mice compared to young control mice, with the decrease particularly pronounced in the old EOC group (Figure S3P). This suggests a diminished pro-inflammatory, classically activated macrophage response in the presence of EOC, especially with aging. Interestingly, despite this decline, the levels of CD38+iNOS+ macrophages were significantly elevated in old EOC mice compared to their age-matched non-tumor controls, indicating a complex regulation of M1 macrophage subsets within the aged tumor microenvironment. On the other hand, the intracellular M2 marker Arginase 1 (EGR2+Arg1+) did not show significant changes across the groups (Figure S3Q). These findings suggest that aging impacts macrophage polarization more in tumor-bearing mice, with older mice showing a shift towards a more immunosuppressive M2 phenotype. While macrophage numbers are not significantly reduced in either young or old tumor-bearing mice, the M1/M2 balance shifts in older mice, which may contribute to a less effective immune response against EOC. This data suggests that compromised macrophage function, along with increased immunosuppressive Tregs, exacerbate the impaired anti-immune response in old mice, potentially contributing to rapid progression of EOC in older mice.

### Aging diminishes the intratumor T cell response in EOC

To validate that the age associated dampened systemic antitumor immune response was mirrored in tumors and TME, we characterized major T cell subsets in the tumors and ascites of young and old mice with ID8^p53-/-^ and ID8^p53-/-, BRCA^ ^1-/-^ tumors. ID8^p53-/-^ tumors of the old mice showed significant decrease in the overall number of CD8^+^T cells (Figure 2A-D), with a marked reduction in the intracellular expression of key effector and cytotoxic markers, including IFNγ, Grz b+, and perforin (Figure 2E-G), while the exhaustion marker PD-1 was increased (Figure 2H) compared to young EOC mice. Similarly, a significant decrease in overall CD4^+^T cells (Figure 2I,J) along with reduction in effector CD4+IFNγ+ and CD4+Tbet+ cells (Figure 2K,L) was observed in tumors from old mice compared to young mice. Most impressively, tumors from old mice demonstrated a significant increase in the overall number of Tregs (Figure 2M-O), and Tregs expressing suppressive cytokines IL-10 and TGFβ (Figure 2P,Q), while PD1 expression was unchanged compared to tumors from young mice (Figure 2R). An identical pattern was observed in the ascites, where we noted a decrease in both CD4^+^ and CD8^+^T cell frequency and the intracellular effector markers (Figure S4A-L), with a concurrent increase in Tregs frequency and IL-10 and TGFβ (Figure S4M-Q). Unlike the intratumoral Tregs, we observed increased PD1 expression in Tregs from ascites (Figure S4R). These findings point to an enhanced immunosuppressive Treg response in the tumors and ascites of old mice with EOC, which may further hinder effective antitumor immunity.

**Figure 2:**
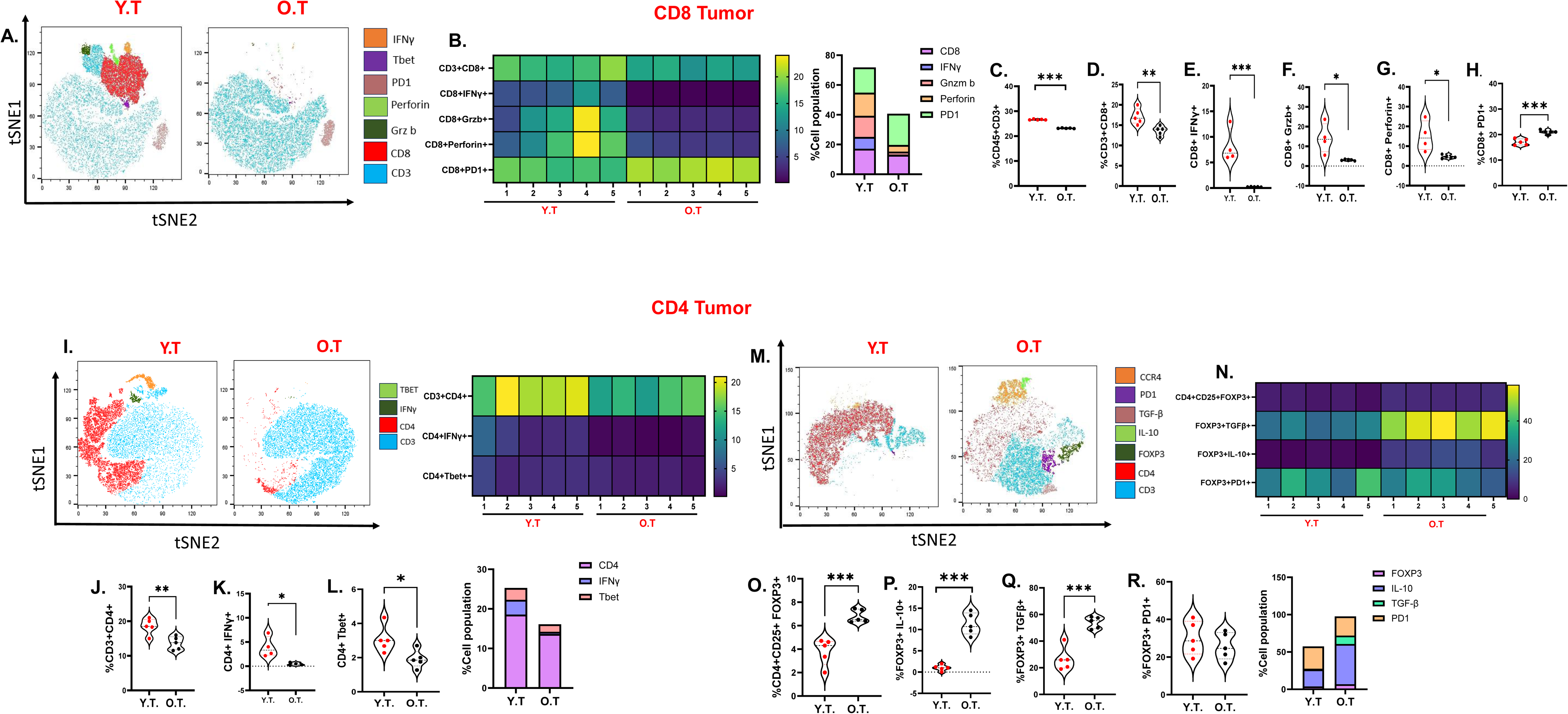
Aging diminishes the intra-tumor T cell response in EOC: ID8^p53−/−^ EOC injected in Y.T or O.T (n = 10/group), Immune profiling was performed in tumors of 5 individual mice per group (A–R) (A) A representative t-SNE visualization of markers after gating on single, live, CD45^+^ CD3^+^, CD4^+^, CD8^+^, IFNγ, Granzyme B, perforin and PD1 expression. (B) Heatmap represents marker expression of the main T cell subsets in individual samples. Violin plots represent the percentage of T cell subsets (C) CD45+CD3+ (D) CD8+, (E) CD8+IFNγ+, (F) CD8+GrzB+, (G) CD8+perforin, (H) CD8+ PD1. (I) A representative t-SNE visualization of markers after gating on single, live, CD45^+^ CD3^+^, CD4^+^, IFNγ, and Tbet expression. Violin plots represent (J) CD4+, (K) CD4^+^ IFNγ+ (L) CD4+Tbet. (M) A representative t-SNE visualization of markers after gating on single, live, CD45^+^ CD3^+^, CD4^+^, CD25+, FOXP3, IL-10, TGF-β and PD1 expression. (N) Heatmap represents marker expression of the main T cell subsets in individual samples. Violin plots represent the percentage of T cell subsets (O) FOXP3+, (P) FOXP3+ IL-10, (Q) FOXP3+ TGFβ, (R) FOXP3+PD1 in the tumors. The experiment was repeated twice in two different sets of mouse experiments. ∗p < 0.05, ∗∗p < 0.01, ∗∗∗p < 0.001, OT compared to YT group by Student’s *t* test.

Tumors from young and aged mice with ID8^p53-/-, BRCA1−/−^ tumors showed a slightly different profile where no significant changes in the overall percentage of CD4^+^T cells or CD8^+^T cells (Figure S5A,B,H) between young and aged mice were found. However, a marked reduction in key effector T cell populations including CD4+IFNγ (Figure S5C), CD4+T-bet+ (Figure S5D), CD8+IFNγ+ (Figure S5I), CD8+Grz b+ (Figure S5J), and CD8+perforin+ (Figure S5K) was observed. Thus, while infiltrating T cell numbers were unchanged, their antitumor function was significantly impaired. Consistently, we observed a significant increase in the proportion of Tregs in ID8^p53-/-, BRCA^ ^1-/-^ tumors of old mice (Figure S5E), with elevated intracellular markers IL-10 and TGFβ (Figure S5F,G). Overall, our findings demonstrate that aging in preclinical models of EOC is associated with an increase in immunosuppressive Tregs and a reduced antitumor T cell response.

### Tregs from old mice exhibit increased immunosuppressive ability

Since the most significant changes were observed in Tregs, which inhibit T cell proliferation and effector activity (M. L. Chen et al., 2005; Schmidt, Oberle, & Krammer, 2012), we assessed the immunosuppressive ability of Tregs from old and young mice. We co-cultured splenic Tregs isolated from young and old EOC mice with CFSE labeled naïve CD4^+^ and CD8^+^T cells from naïve young mice at a 1:2 ratio in the presence of IL-2 (20 ng/ml) for 72 hours. Flow cytometric analysis showed that Tregs from old EOC mice inhibited the proliferation of both CD4^+^ and CD8^+^T cells to a greater extent compared to Tregs from young EOC mice (Figure 3A,B,D,E) and were particularly effective in inhibiting the production of IFNγ by both CD4^+^ and CD8^+^T cells (Figure 3C,F). In the same co-culture experiment, we profiled key senescence markers on CD4^+^ and CD8^+^ T cells and observed a significant reduction in CD28 percentage(Figure 3I,J,N), a co-stimulatory molecule essential for T cell activation (Effros, 1997), in the presence of Tregs derived from old EOC mice compared to young EOC mice. Additionally, CD57, a marker associated with terminally differentiated or senescent T cells with diminished proliferative capacity (Cura Daball et al., 2018), was significantly increased in both CD4^+^ and CD8^+^T cells from aged EOC mice (Figure 3K,O,R). Other senescence markers, such as KLRG1 (Figure 3L,P,R) and β-galactosidase (Figure 3M,Q,R), which reflect a permanent loss of proliferative potential (X. Liu et al., 2018; Ramello et al., 2021), were also significantly elevated in CD4^+^ and CD8^+^T cells from aged EOC mice. Next to see if EOC TME can influence Tregs, we conducted a Click-IT EdU proliferation assay on naive Tregs exposed to ascites isolated from EOC-bearing young and old mice. Tregs exposed to ascites from old mice underwent increased proliferation, shown by increased EdU incorporation, compared to Tregs cultured in ascites from young mice (Figure 3G,H). Together, these observations show strong evidence that Tregs from old mice have enhanced suppressive function and promote T-cell senescence as a suppressive mechanism (X. Liu et al., 2021; J. Ye et al., 2012), contributing to an increased immunosuppressive TME and potentially facilitate immune evasion by EOC.

**Figure 3:**
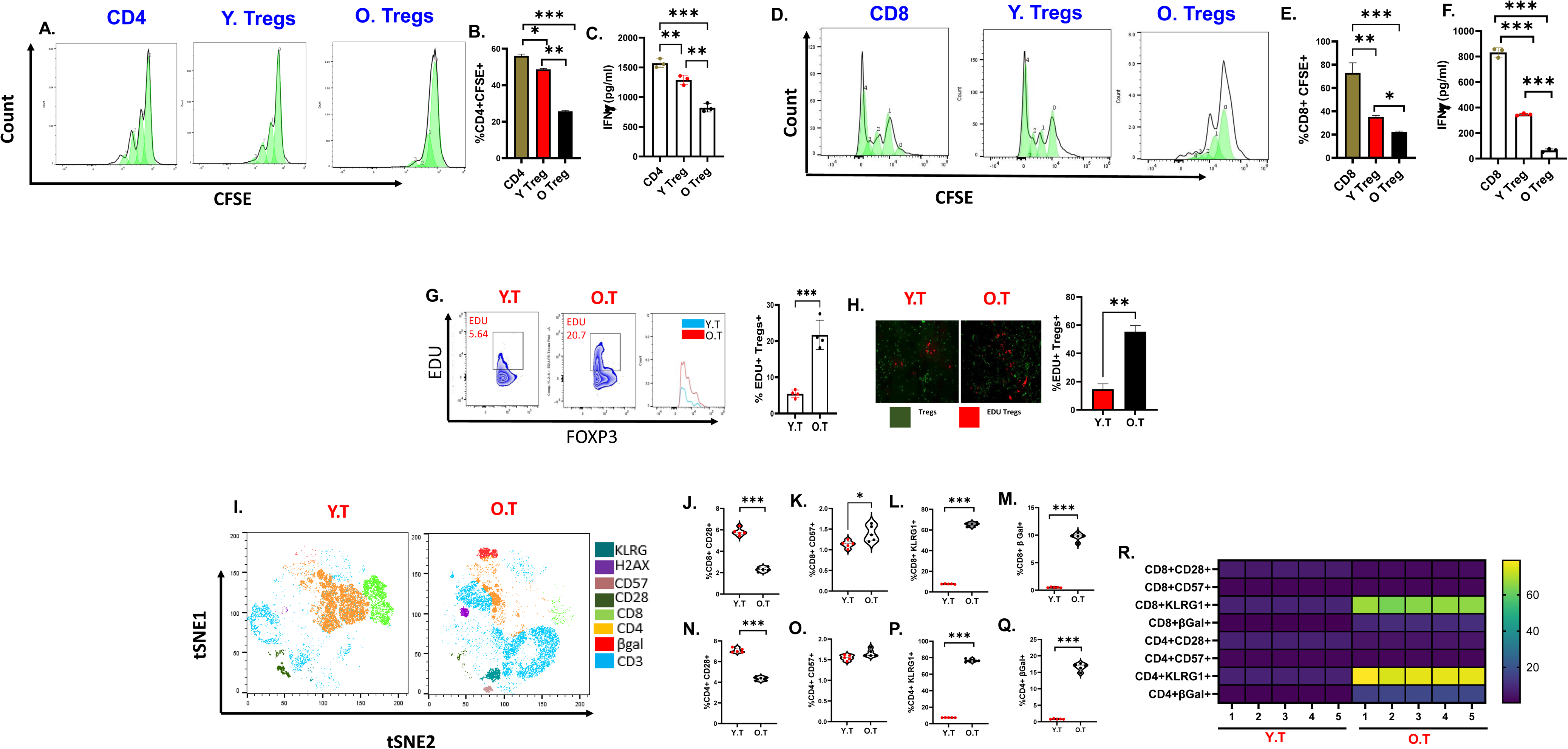
Tregs from old mice are more immunosuppressive: Naïve CD4 and CD8 T cells were isolated and were labelled with CFSE for 20 mins. These cells were co cultured with Tregs from young and old EOC mice at a ratio of 2:1. (A) CFSE proliferation histograms representing CD4+ proliferation in presence of Tregs from old and young EOC mice. (B) Bar graph represents %CD4+CFSE. (C) Bar graph represents IFNγ levels from CD4: Treg co-culture experiment. (D) CFSE proliferation histograms representing CD8 proliferation in presence of Tregs from old and young EOC mice. (E) Bar graph represents %CD8+CFSE. (F) Bar graph represents IFNγ levels from CD8: Treg co-culture experiment. (G) Flow plots and bar graph representing %FOXP3+ EDU. (H) Immunocytochemistry images representing EDU+Tregs in red. (I) A representative t-SNE visualization of markers after gating on single, live, CD45^+^ CD3^+^, CD4^+^, CD8+, CD28, CD57, H2AX, βGal and KLRG. Violin plots represent the percentage of senescence markers on T cell subsets (J) CD8+ CD28, (K) CD8+CD57, (L) CD8+ KLRG1 and (M) CD8+ βGal. (N) CD4+CD28, (O) CD4+ CD57, (P) CD4+ KLRG1 and (Q) CD4+βGal. (R) Heatmap represents marker expression of the main senescence markers on T cell subsets in individual samples. ∗p < 0.05, ∗∗p < 0.01, ∗∗∗p < 0.001, OT compared to YT group by Student’s *t* test.

### Tregs are key in promoting EOC in aged mice

To investigate if Tregs are crucial in promoting EOC progression in old mice, we utilized the anti-CD25 (clone PC-61) antibody to deplete Tregs in ID8^p53-/-^ tumor-bearing old mice (Figure S6A) (Kalim et al., 2022; Y. Li et al., 2023). Depletion of Tregs significantly enhanced the survival of the old mice, with an increase in median survival to 64 days, compared to 45 days in the control antibody group (Figure S6B). The decreased tumor burden was reflected by low ascites accumulation in the depletion mice compared to control mice (Figure S6C). Enhanced immune response was seen in the Treg-depleted mice as indicated by significant increase in CD4^+^ and CD8^+^T cell populations in the peripheral blood and ascites compared to control mice (Figure S6E,K). Treg depletion also enhanced levels of effector CD4+IFNγ+ and CD8+IFNγ+ (Figure S6F,L) cells. Interestingly, we observed a trend in increased perforin expression (Figure S6M), while Grz b+ levels were unchanged (Figure S6N) between the depletion and control groups. Surprisingly, Treg depletion also reduced Treg immunosuppressive markers like FOXP3+, IL-10, and TGFβ (Figure S6G-I) (Jarnicki, Lysaght, Todryk, & Mills, 2006). Treg depletion played a role in modulating T cell senescence as reflected by a significant decrease in the percentages of CD8+β Gal+, CD4+β Gal+, and CD4+CD57+ (Figure S6Q,U,V) cells compared to control mice. However, no significant differences were found in the percentages of CD8+KLRG1+, CD8+CD57+, CD8+CD28+, or CD4+KLRG1+ (Figure S6P,R-T) cells between control and Treg-depleted aged mice. Notably, Treg depletion led to an increase in the percentage of CD4+CD28+ (Figure S6W) cells. Together, these findings underscore the pivotal role of Tregs in promoting EOC progression in aged mice, and absence of Tregs allows for a more robust immune response against EOC cells.

### Tregs from aged EOC mice prefer OXPHOS for immunosuppression

T cell function, differentiation, and activation are largely governed by metabolic modifications (Kouidhi, Elgaaied, & Chouaib, 2017; Ma, Ming, Wu, & Cui, 2024). Previous research has shown Tregs to be dependent on OXPHOS for activation and function (A. Han et al., 2023; Yan et al., 2022). Bioenergetics of splenic Tregs isolated from young and old mice with and without EOC were profiled by XFe Seahorse analyzer. At baseline level, the Tregs from young mice without EOC displayed lower OXPHOS measured as OCR, which did not change even under stress. The Tregs from old mice with EOC showed significantly increased OXPHOS compared to Tregs from young tumor-bearing mice, and both young and aged healthy controls, under both basal and stressed conditions (Figure 4A,B,E and S7A,B). The basal ECAR, which reflects glycolytic activity, did not differ between young and old control or tumor-bearing Tregs at basal level; however, old Tregs from EOC mice exhibited increased glycolysis under stressed conditions when compared to young Tregs from EOC mice and their controls (Figure 4C,D and S7C,D). The resulting energy phenotype indicated a basal aerobic energy phenotype in the Tregs from old EOC mice that further moved to an energetic phenotype when stressed (Figure 4E). To validate the observations of splenic Tregs, we used SCENITH to profile the metabolic dependency of Tregs from ascites of young and old mice. Tregs from old EOC mice had higher mitochondrial dependency, decreased glycolytic capacity and non-significant increase in Fatty acid oxidation capability compared to Tregs from young EOC mice (Figure 4F-H). While no significant changes were observed in glucose dependency between young and old Tregs (Figure 4I). When Tregs were treated with OXPHOS inhibitor oligomycin, Tregs from old mice showed a decrease in expression of key immunosuppressive markers, including FOXP3, TGFβ and IL-10 (Figure 4K-M), while treatment with the glycolysis inhibitor 2DG had no effect. Oligomycin treatment resulted in reversal of CD4^+^T cell suppression, indicating a functional reprogramming of Treg cells (Figure 4M). This effect was observed in comparison to treatment with 2DG, which primarily targets glycolysis (Figure 4M). We also observed that inhibiting glycolysis led to a significant decrease in IFNγ production, Importantly, treatment with oligomycin reversed the suppression of IFNγ levels, further supporting the idea that glycolysis plays a crucial role in maintaining the effector functions of T cells (Figure 4O). These findings suggest that inhibiting OXPHOS in Tregs from aged mice not only reduces their suppressive capacity but also leads to an enhancement of CD4^+^ T cell responses, potentially re-establishing more effective antitumor immunity.

**Figure 4:**
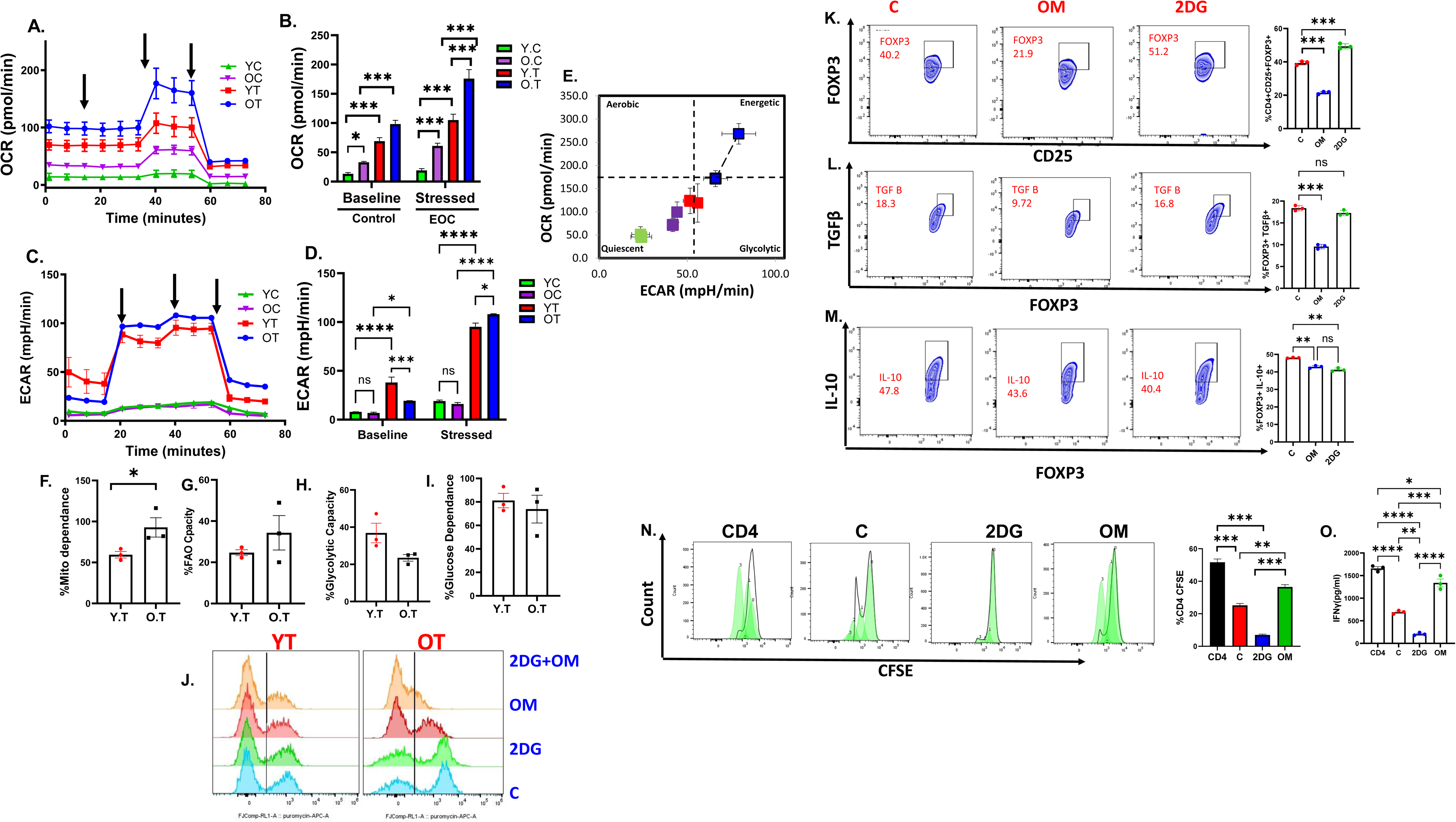
Tregs from aged EOC mice prefer OXPHOS for immunosuppression: ID8^p53−/−^ EOC injected in Y.T or O.T, FOXP3+ cells were isolated from spleens of control and EOC young and old tumor at 5 weeks and subjected to Seahorse analysis and energy targeted metabolomics. (A) Oxygen consumption rate (OCR) was assessed in real-time using an XFe Seahorse analyzer as described in methods. Port injections of (1) oligomycin, (2) FCCP, and a combination of (3) rotenone-antimycin were given. (B) The bar graph represents basal and stressed OCR (n = 3). (C) Extracellular acidification rate (ECAR) was measured with port injections of (1) glucose, (2) oligomycin, and (3) 2-DG. (D) The bar graph represents basal and stressed ECAR (n = 3). (E) A quadrant plot indicates the metabolic shift of energy phenotype at basal and stressed levels. SCENITH was performed in Tregs isolated from young and old EOC mice. Bar graphs represent the (F) %Mitochondrial dependence, (G) % FAO Capacity, (H) %Glucose dependance, (I) Glycolytic capacity. (J) Histogram representing the FOXP3+ Puromycin in Tregs isolated from young and old EOC mice after treatment. Tregs isolated from young tumor bearing mice were treated with 2DG and oligomycin, (K) Flow plots and bar graph represents FOXP3 percentage. (L) Flow plots and bar graph represents TGFβ percentage. (M) Flow plots and bar graph represents IL-10 percentage. (N) CFSE proliferation histograms and bar graph representing CD4 proliferation in presence of Tregs treated with 2DG and oligomycin. (O) Bar graph represents IFNγ levels from CD4: Treg co-culture treated with 2DG and oligomycin. ^∗^p < 0.05, ^∗∗^p < 0.01, ^∗∗∗^p < 0.01 tumor compared to control as assessed by unpaired T-test.

### Succinate as a metabolic regulator of Tregs in aged mice with EOC

To gain an understanding of the metabolic alterations, we performed a targeted metabolomics in the Tregs by liquid chromatography-tandem mass spectroscopy (Udumula et al., 2024; Udumula et al., 2023). Analysis of glycolytic metabolites in Tregs revealed no significant differences in glucose (Figure S8A), fructose (Figure S8B), galactitol (Figure S8C), or glucose-6-phosphate (Figure S8E) between young and old control or EOC-bearing mice. However, glycerate levels were elevated in Tregs from old EOC mice compared to their young counterparts (Figure S8G). Additionally, phosphoenolpyruvate (Figure S8D) and mannitol (Figure S8F) levels were increased in Tregs from old EOC mice compared to both young EOC mice and old controls. A striking observation was a strong increase in succinate in Tregs from old mice with EOC compared to young EOC mice, as well as in old non-EOC mice compared to young non-EOC mice (Figure 5D). αketoglutarate (αKG) was significantly increased in old control Tregs while it decreased in EOC conditions (Figure 5C), which may indicate a rapid conversion of αKG into succinate. Other downstream metabolites such as fumarate, malate, isocitric acid, and glutamate (Figure 5B,E,F,H) were significantly decreased in old Tregs compared to young Tregs from EOC mice. Citric acid levels decreased in old tumor Tregs when compared to old control Tregs; however, there was no significant difference between young and old tumor Tregs (Figure 5A). Lactate levels were also significantly decreased in older Tregs when compared to other groups (Figure 5I). Pyruvic acid levels were not changed between any of the groups (Figure 5J). To support the increased succinate levels, we measured succinate dehydrogenase levels using a SDH colorimetric assay in Tregs isolated from young and older EOC mice and observed decreased SDH levels in Tregs from old EOC mice compared to young mice, which may suggest that succinate is not getting converted to fumarate and agrees with the accumulation of succinate (Figure 5K-N). The succinate receptor SUCNR1 (also known as GPR91) was also significantly increased in older Tregs when compared to younger EOC Tregs and healthy controls (Figure 5O). Cancer cells release multiple soluble factors that not only stimulate their growth and metastasis but also regulate immune cells in the TME to facilitate their progression (Cheng et al., 2023; Cortellino & Longo, 2023; Mortazavi Farsani & Verma, 2023), an interplay that is crucial for tumor initiation and progression. To investigate whether EOC in older mice contributes to increased succinate levels, we performed colorimetric succinate measurement, which showed an increase in succinate in ascites and tumor cells (Figure 5P,Q). As further validation, targeted metabolomics in serum of aged (≥ 65 years) and younger (≤ 55 years) EOC patients and matched healthy controls corroborated that aged EOC patients exhibited significantly increased succinate levels compared to both younger EOC patients and healthy young and old controls (Figure 5R,S), validating increased succinate being produced by the tumor or TME. This data demonstrates that along with Tregs, the aged TME and tumors have increased succinate levels, which may be taken up by the Tregs.

**Figure 5:**
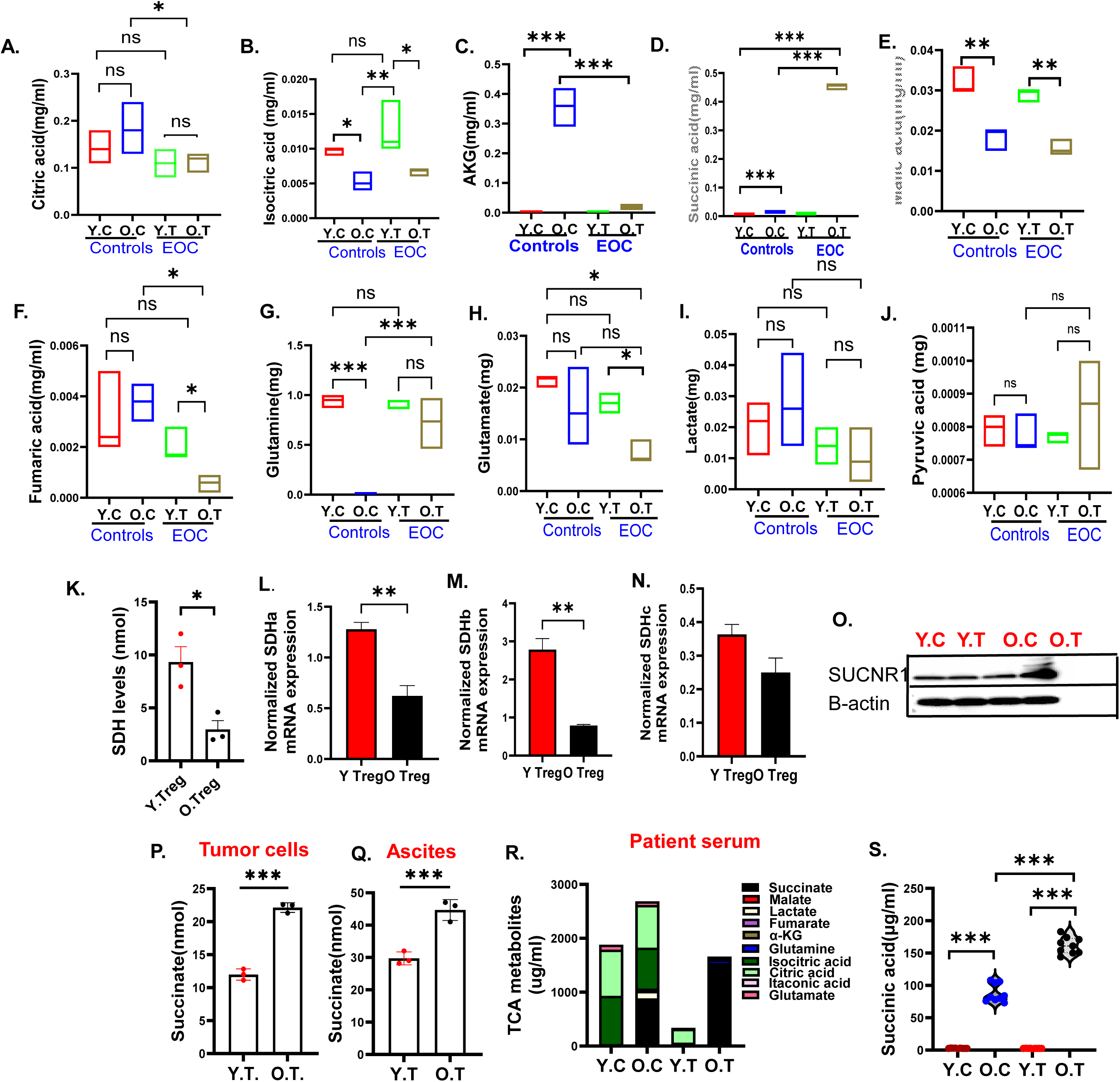
Succinate as a metabolic regulator of Tregs of aged mice with EOC: Tregs were isolated from control and tumor-bearing young and old mice and were processed for Targeted TCA analysis. (A-J) Targeted analysis of the TCA cycle metabolites was performed to assess the levels of various metabolites in pooled xenografts (*n* = 3) in triplicates. (K) Bar graph represents SDH levels in Tregs isolated from young and old EOC mice. Normalized mRNA expression of (L) SDHa, (M) SDHb, (N) SDHc in Tregs of young and old EOC mice. (O) Immunoblot representing SUCNR1 expression in Tregs of young and old control and EOC mice. (P) Bar graph represents succinate levels in tumor cells of young and old EOC mice. (Q) Bar graph represents succinate levels in ascites of young and old EOC mice. (R) Bar graph represents levels of all TCA metabolites in young and old health and ovarian patients’ serum. (S) Violin plot represents individual values of succinate in young and old health and ovarian patients’ serum from R. ^∗^p < 0.05, ^∗∗^p < 0.01, ^∗∗∗^p < 0.01 tumor compared to control as assessed by unpaired T-test.

### Tumor derived succinate promotes Treg function in aged EOC mice

To investigate the role of succinate in regulating Treg function, splenic Tregs isolated from young EOC mice were treated with varying concentrations of succinate for 48 hours (100 μM, 1 mM, and 5 mM). Succinate resulted in a significant increase in the expression of CD25+FOXP3 and other immunosuppressive markers, IL-10 and TGFβ (Figure S9A-C). The addition of succinate to the co-culture of CD4^+^T cells and Tregs reduced the proliferation of CD4^+^T cells (Figure S9D), confirming succinate’s crucial role in modulating Treg function. To explore whether succinate from the TME can enhance Treg activity, Tregs from naive mice were exposed to serum from both young and old patients, where serum from old patients led to an increase in FOXP3 expression and immunosuppressive markers IL-10 and TGFβ (Figure S9E-G), demonstrating that systemic succinate enhances Tregs function. To investigate the impact of tumor-derived succinate on Tregs, succinate synthesis in tumor cells derived from aged EOC mice was inhibited using CPI-613, a compound that blocks α-ketoglutarate dehydrogenase complex (αKGDC), mediated succinate production from αKG (Gao et al., 2020; Khan et al., 2023; Udumula et al., 2024). CPI-613 led to a reduction in succinate levels in tumor cells compared to untreated cells (Figure S9H). The tumor-conditioned media (TCM) collected from CPI-613 treated tumor cells (TCM+CPI-613) was then added to Treg cultures, which resulted in a reduction in Tregs, decreased expression of FOXP3, IL-10 and TGFβ (Figure S9H-K). In addition, the low succinate TCM reversed CD4^+^ T cell suppression compared to TCM from untreated tumor cells (Figure S9L). Moreover, key senescence markers of CD28 was increased and KLRG1 was significantly reduced in the Treg-CD4 co-culture exposed to TCM+CPI-613, while no significant changes were observed in CD57 and β-galactosidase (Figure S9M-P). Further Tregs exposed to TCM+CPI-613 displayed decreased OXPHOS when compared to TCM exposed Tregs (Figure S9Q,R), while we did not observe any differences in ECAR between TCM and TCM+CPI-163 exposed Tregs (Figure S9S-U). These findings collectively support the conclusion that succinate derived from tumor cells or the TME plays a significant role in enhancing Treg function.

### αKGDC inhibition improves overall survival in old EOC mice

AA6 is a selective inhibitor of the αKGDC, which blocks the conversion of αKG to succinate (Atlante et al., 2018). We evaluated the effect of AA6 on Tregs, the TCM collected from AA6 treated tumor cells (TCM+AA6) was then added to Treg cultures, which resulted in a reduction in Tregs, decreased expression of FOXP3, IL-10, TGFβ and PD1 (Figure S10A-H). Further we administered AA6 to ID8^p53-/-^ EOC-bearing young and old mice. AA6 treatment significantly prolonged survival in old EOC mice compared to untreated mice, while no survival benefit was observed in young EOC mice (Figure 6A). Correspondingly, tumor burden, assessed by ascites volume, was significantly reduced in older AA6 treated mice (Figure 6B), while no difference was detected in the young mice. Treatment with AA6 led to a reduction in succinate levels within the ascitic fluid of both young and old mice with ID8^p53-/-^ tumors (Figure 6C). Thus, this could be attributed to low succinate levels in young EOC mice, suggesting a limited therapeutic window for AA6 in this group. Immune profiling revealed an increase in the percentage of CD4^+^ and CD8^+^T (Figure 6D,E,J) cells in both young and old AA6 treated mice relative to their respective controls. However, enhanced effector function, as observed by increased levels of CD4+IFNγ+, CD8+IFNγ+, and CD8+granzyme B+ (Figure 6F, K, L) T cells, was observed only in aged AA6-treated mice, while these were unchanged in the young EOC mice. AA6 treatment in EOC old mice increased CD8+PD1+ percentage in old EOC mice, while it was unchanged in young EOC mice (Figure 6M). More importantly, the frequency of Tregs (Figure 6G) was reduced in both young and old AA6-treated EOC mice compared to their respective untreated controls. In aged mice, AA6 treatment resulted in decreased immunosuppressive Treg subsets expressing IL10 and TGFβ (Figure 6H,I), an effect not observed in the young group. AA6 treatment also led to a decrease in markers of T cell senescence like CD57, KLRG1, and β-galactosidase (Figure 6N-P) on CD8^+^T in both young and old AA6-treated mice compared to untreated controls. A similar decline was observed in CD4^+^T (Figure 6Q-S) cells expressing these senescence markers across both age groups. Taken together, these findings demonstrate that AA6 exerts age-dependent antitumor effects by modulating metabolic pathways, reducing immunosuppression, and attenuating T cell senescence. The therapeutic benefit observed predominantly in aged mice underscores the importance of considering host age and metabolic context when developing targeted therapies for EOC.

**Figure 6:**
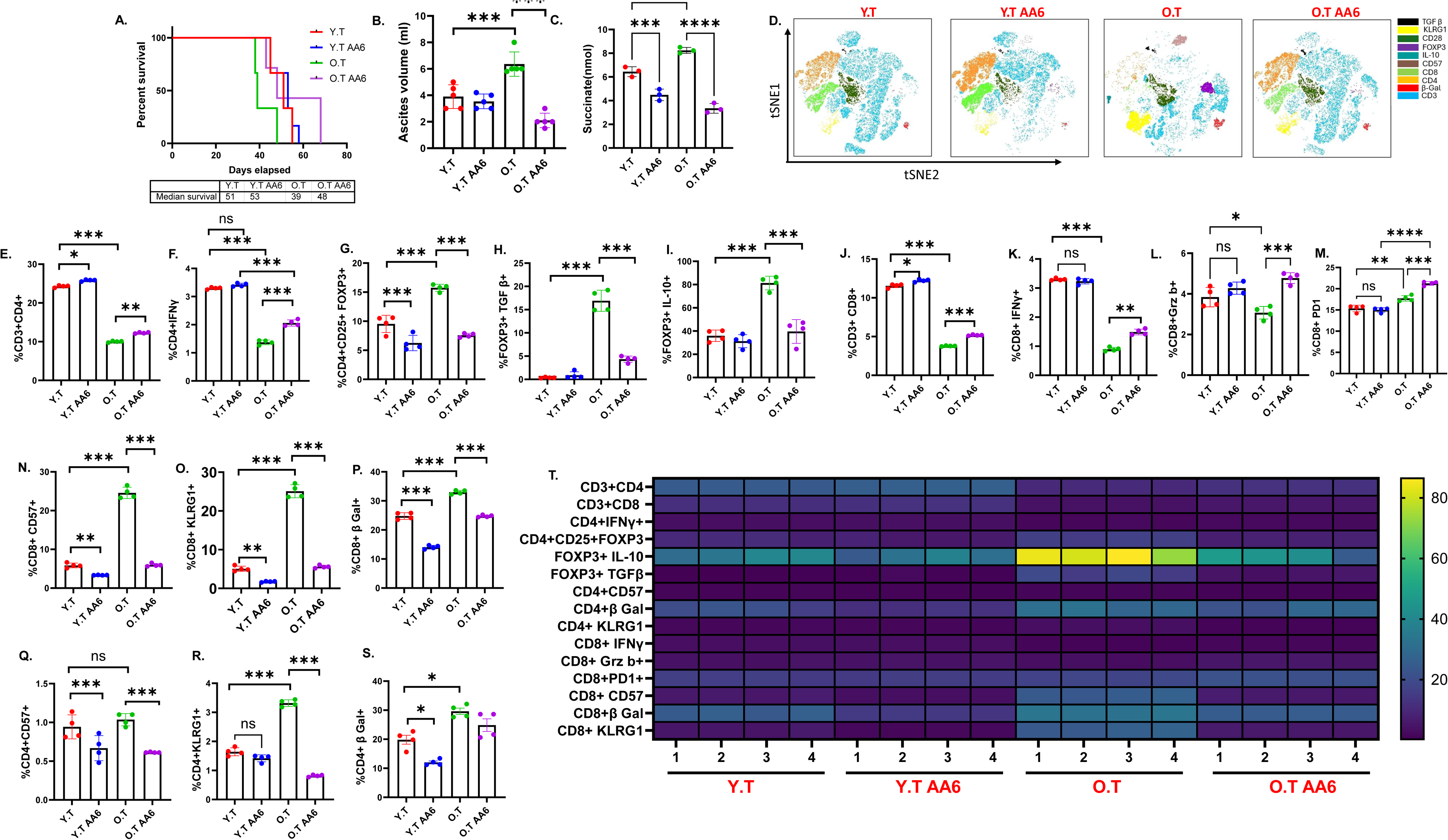
aKGDC inhibition improves overall survival in old EOC mice: Kaplan–Meier graphs indicating overall survival in mice bearing (A) ID8^p53-/-^ cells were injected in mice and were randomly assigned for Y.T, Y.T AA6, O.T, O.T AA6 groups. (B) Bar graphs represent average ascites accumulated in Y.T, Y.T AA6, O.T, O.T AA6 mice. (C) Bar graphs represent succinate levels in ascites of Y.T, Y.T AA6, O.T, O.T AA6 mice. (D) A representative t-SNE visualization of markers after gating on single, live, CD45, CD4, CD8, IFNγ, FOXP3, IL-10, TGFβ, CD28, KLRG1, β-Gal. (E) CD4+, (F) CD4^+^ IFNγ+, (G) FOXP3, (H) FOXP3+ TGFβ, (I) FOXP3+ IL-10, (J) CD8^+^, (K) CD8+IFNγ+, (L) CD8+GrzB+, (M) CD8+ PD1, (N) CD8+ CD57, (O) CD8+ KLRG1, (P) CD8+ βGal, (Q) CD4+CD57, (R) CD4+ KLRG1, (S) CD4+βGal. (T) Heatmap represents marker expression of the main T cell subsets in individual samples. Immune profiling was performed in 4 individual mice per group. ∗p < 0.05, ∗∗p < 0.01, ∗∗∗p < 0.001, by one-way ANOVA, followed by Sidak multiple comparison test.

## Discussion

Aging is a primary risk factor for developing EOC and other cancers (Foster et al., 2011; Harman, 1991; Sedrak & Cohen, 2023; Sitnikova et al., 2023). While the average age of EOC diagnosis is 63 years, nearly 45% of women are over 64 and with 25% above 74 years when diagnosed. Unfortunately, EOC outcomes also worsen with age and have poor prognoses (Hightower et al., 1994; Thigpen et al., 1993; Tortorella et al., 2017). Despite this factual relationship between age and EOC, there is limited knowledge about underlying mechanisms. In our study, we demonstrate for the first time, the impaired antitumor T-cell responses in aged preclinical EOC models, driven by overabundance of immunosuppressive Tregs considering progressive decline of the adaptive immune system is an established hallmark of aging, a better understanding of the intersection between immune aging and tumor progression has widespread implications for human health (Hakim, Flomerfelt, Boyiadzis, & Gress, 2004).

Studies have shown that highly immunogenic mouse tumors, such as colon (MC38) and breast (EO771) grow more rapidly in aged animals, while less immunogenic tumors, such as melanoma (B16), do not (Georgiev et al., 2024). EOC is considered a less immunogenic and cold tumor (Blanc-Durand, Clemence Wei Xian, & Tan, 2023; Ghisoni, Imbimbo, Zimmermann, & Valabrega, 2019), but reports have demonstrated EOC to progress aggressively with increased metastasis in aged mice (Chan et al., 2006; Hou et al., 2022). Our study corroborates these reports and shows decreased survival and increased tumor burden in ID8 set of EOC cells lines. Aging is associated with a gradual and sustained rise in systemic pro-inflammatory stress (Isola et al., 2024), often referred to as inflammaging (Di Giosia et al., 2022; Franceschi et al., 2000), and exhibition of a SASP phenotype (Coleman et al., 2013). We found that in EOC-bearing old mice the tumor environment represented by ascites had higher levels of SASP cytokines, indicating a close interaction of the EOC and the aged host environment, which encourages tumor growth. In our preclinical models of EOC we demonstrate that aged mice had decreased survival with increased tumor burden when compared to younger mice afflicted with EOC. Interestingly, the modulation of these factors and cytokines differed between the cell line models, indicating tumor phenotype specificity. For example, IFNγ was increased in the ascites of old mice with ID8^p53-/-, BRCA1-/-^ tumors, but decreased in mice with ID8^p53-/-^ tumors, which is due to a more immunosuppressive tumor microenvironment in aged hosts lacking BRCA1 mutation.

Although age associated inflammatory environment and immune decline is well documented (Falvo et al., 2025; Garg et al., 2014), limited studies have investigated the role of an aged immune system in EOC. A study reported that an age-related increase in metastatic tumor burden was associated with alterations in tumor infiltrating lymphocytes and B cell-related pathways within the gonadal adipose tissue of ovarian tumors (Loughran et al., 2018). However, no studies have profiled the differential immune response in young and old EOC. Studies detailing the change in age related distribution and functionality of T cell subsets shift have demonstrated an increase in Tregs with higher immunosuppressive ability in older individuals as a major characteristic (A. C. Y. Chen et al., 2024; Elyahu et al., 2019; Garg et al., 2014; Han, Georgiev, Ringel, Sharpe, & Haigis, 2023; B. Zhao et al., 2023). While the role of Tregs has been demonstrated (Erfani et al., 2014; Ohue & Nishikawa, 2019; Zhang et al., 2015) in EOC, it is less explored in aging. Our study shows that old EOC mice have a weaker immune response, marked by reduced antitumor CD4⁺ and CD8⁺T cells and increased immunosuppressive Tregs with enhanced suppressive activity, compared to young EOC mice. These observations are similar to studies (Georgiev et al., 2024) in aging models of melanoma and colon cancer, which demonstrate a role for CD8^+^ T cells and a potential role of Tregs. This correlation suggests a role for Tregs in suppression of antitumor T cell response and weakening of the adaptive immune response in old mice with EOC. This was supported by the observations that Tregs from old mice with EOC expressed increased immunosuppressive cytokines such as TGFβ and IL-10.

Furthermore, the Tregs from old mice with EOC were able to suppress proliferation and IFNγ production in both CD4^+^ and CD8^+^T cells. Importantly, depletion of Tregs in old mice improved survival and reduced tumor growth in both young and old mice, with a more pronounced effect in old mice with EOC. The improved outcome in old mice was marked by an increase in CD4^+^T cells, CD8^+^ T cells and the upregulation of effector cells, specifically CD4+IFNγ and CD8+IFNγ and non-significant upward trend in granzyme b percentage. Similarly, Treg depletion impacted only β-Gal on CD4 and CD8 T cells and CD57 and CD28 on CD4^+^T cells, while KLRG1 levels were unchanged, but PD1 was significantly decreased. Our results suggest that Treg depletion may have a selective impact, affecting only certain effector-related and senescence-related markers. This finding contrasts with other studies, which demonstrate Tregs to suppress T cells by inducing responder T-cell senescence (X. Liu et al., 2018; J. Ye et al., 2012). These findings are consistent with other published studies, which have shown that Treg depletion enhances the effectiveness of Immune check point therapies in reducing tumors in melanoma and other cancers (Curtin et al., 2008).

Metabolites play a crucial role in the proper functioning of immune and cancer cells. Tregs engage nutrient sensing mechanisms to adapt to both intrinsic and extrinsic environmental cues, triggering metabolic reprogramming to sustain their activity (Kempkes, Joosten, Koenen, & He, 2019). Tregs have been demonstrated to rely predominantly on OXPHOS and FAO, to support their suppressive functions (Field et al., 2020; X. Zhao et al., 2025), although glycolysis has also been shown to be crucial for their migratory bioenergetic demands. Recent non-cancer studies have shown dependence of aging Tregs on deactivating oxidative stress while maintaining high OXPHOS and enabling them to function, grow, and activate in older mice (Danileviciute et al., 2022; Guo et al., 2020). Research on aging-related metabolic changes in Tregs in EOC is limited. Our study is the first to show that Tregs from old EOC mice have higher OXPHOS and ECAR levels than those from young or non-EOC mice, suggesting greater metabolic fitness with age and cancer. SCENITH analysis showed these Tregs rely on mitochondria, not glycolysis or fatty acid oxidation. Blocking OXPHOS reduced FOXP3 and TGFβ, but not IL-10, and restored CD4⁺T cell function. In contrast, blocking glycolysis had no effect on Tregs but impaired CD4⁺T cells, showing their dependence on glycolysis for activity (Chang et al., 2013; S. Liu et al., 2023).

The plasticity of Treg metabolism and their ability to utilize tumor/TME metabolites to regulate behavior and function has been well-documented in multiple cancers (Gu et al., 2022; Moon, Hajjar, Hwu, & Naing, 2015; Phan & Goldrath, 2015; Shi et al., 2011; Wang, Franco, & Ho, 2017). However, no age specific metabolite in regulation of Tregs in EOC, especially in aged EOC has been defined. Targeted metabolomics showed that succinate levels in Tregs were higher in old control mice than in young control mice, and even higher in old EOC mice compared to young EOC mice. Succinate, a key metabolite in the TCA cycle, has been implicated in the regulation of immune cell function, including Tregs (Kinsella et al., 2023). SDH enzyme, which converts succinate to fumarate, levels were low in older Tregs when compared to younger Tregs. Further, succinate receptor SUCNR1 was also increased in Tregs isolated from old EOC mice. This suggests that Tregs TCA cycle to promote succinate accumulation as well as ability to imbibe extracellular succinate. Although the impact of succinate on Tregs in cancer has not been explored much, studies have shown that succinate uptake in CD4^+^T cells suppress their effector function by inhibiting mitochondrial glucose oxidation (Gudgeon et al., 2022). A recent study elegantly showed extracellular succinate to promote Treg numbers in lung cancer and melanoma cells (Kinsella et al., 2023). Other research has demonstrated that extracellular succinate can polarize macrophages into the M2 phenotype through SUCNR1 signaling (Harber et al., 2020; Trauelsen et al., 2021). In our studies, Tregs isolated from younger mice exposed to succinate resulted in a dose-dependent increase in the expression of FOXP3, TGFβ, and IL10, thereby enhancing their immunosuppressive phenotype, as functionally seen by suppression of CD4^+^T cell proliferation in a dose dependent manner. Upregulated SUCNR1 in Tregs from old mice with EOC indicated increased potential to uptake succinate, which raised the questions if EOC cells and TME are sources of succinate. We found that tumors and ascites from old EOC mice did indeed have elevated succinate. This was further confirmed in patients where serum from EOC patients over the age of 65 demonstrated significantly increased succinate compared to other TCA cycle metabolites, in comparison to EOC patients below 55 years of age. Exposure of the patient serum or ascites from old EOC mice to Tregs from naïve young mice increased the Treg number as well as expression of immunosuppressive markers. As further proof, when succinate production in tumor cells derived from aged mice was inhibited, the conditioned media exposure with low succinate, resulted in a marked reduction of FOXP3 expression and immunosuppressive cytokines TGFβ and IL-10, indicating a decrease in suppressive phenotype of Tregs, confirmed by reversal of CD4^+^T cell proliferation. Succinate inhibition also led to a reversal in the expression of key senescence markers, including CD28 and KLRG1. These results indicate a pivotal role of succinate in regulating immunosuppressive Treg function in an aged TME of EOC. To the relevance of succinate in vivo, we employed a relatively more specific αKGDC inhibitor, AA6. AA6 treatment improved survival and reduced tumor burden in old EOC mice but had no effect on young mice. It decreased FOXP3⁺ cells in both groups, while reducing immunosuppressive IL-10 and TGFβ only in old mice. In old mice, AA6 also restored antitumor immunity by increasing CD4⁺, CD8⁺, and cytotoxic T cells and lowering senescence markers. Inhibition of αKGDC also resulted in increased PD-1 expression on CD8^+^T cells, which may reflect enhanced T cell activation (Simon & Labarriere, 2017). Collectively, our study demonstrates that aged TME of EOC is characterized by elevated succinate, which is key in promoting Treg-mediated immunosuppression probably by ensuring metabolic fitness and contributing to effective immune escape in elderly EOC patients. Targeting succinate metabolism may therefore offer a promising strategy for modulating the immune landscape and restoring antitumor immunity in aging individuals. However, our study has some limitations, as inhibition of aKGDC complex by CPI-613 or AA6 will result in accumulation of αKG, another active metabolite with potential to regulate signaling and epigenetic processes. αKG by itself has been demonstrated to have antitumor (N. Liu et al., 2023; Xiang et al., 2024) effects on cancer cells directly, regulate cytotoxic T cell function and proliferation (Minogue et al., 2023) and is crucial for TET family function of demethylation (Tran, Dillingham, & Sridharan, 2019). The anti-aging properties of αKG, via modulating cellular senescence are also well documented (Asadi Shahmirzadi et al., 2020). Our study does not address the impact of αKG on EOC when succinate production is inhibited. Another point that can be further explored in depth is the succinate source. Although, we demonstrate strongly the role for extracellular succinate, we do not present the importance of intrinsic TCA rewiring that may also lead to increased succinate production.

Despite this, our study implies a crucial role for Tregs in driving aggressive EOC due to increased immunosuppression in an already age-compromised environment. Thus, understanding the changes and mechanisms of Treg dynamics with age can help tailor precision-based therapeutics including immunotherapy for older EOC patients.

## Acknowledgements

The authors thank Ms. Stephanie Stebens at Henry Ford Health for editorial assistance.

## Conflicts of Interest

The authors declare no conflict of interest.

## Funding

This work was supported by NIH/NCI R01CA249188 to R.R. MP.U is supported by Henry Ford Health Internal Mentored Award (A20091). K.M. is partially supported by Henry Ford Cancer post-doctoral fellowship award. S.G. is supported by grants from the National Multiple Sclerosis Society (RG-1807-31964, RG-1508-05912), the NIH (NS112727, AI144004).

## Permissions

### Author Contributions

MP.U. designed the experiments, conducted research, analyzed the data, wrote and edited the manuscript. A.KM., M.N., F.R., H.S., and T.B. performed research and analyzed data. M.H., S.G., EN.C., and H.M.G. analyzed the data and edited manuscript. R.R. designed experiments, supervised the research, analyzed the data and edited the manuscript.

### Data Availability

This study does not analyze any publicly available datasets. All data supporting the findings of this study are available from the corresponding author upon reasonable request. No original code was generated or used in this study. Any additional information necessary to reanalyze the reported data can be obtained from the corresponding author upon request.

---

**Table 1:** List of flow antibodies.

| Reagent | Source | Identifier |
| --- | --- | --- |
| CD45 | S18009F | Biolegend |
| CD3 | 17A2 | Biolegend |
| CD4 | GK.1.5 | Biolegend |
| CD8 | 53–6.7 | Biolegend |
| CD25 | PC61 | Biolegend |
| PD1 | RMP1-30 | Biolegend |
| IFNγ | XMG1.2 | Biolegend |
| IL-10 | JES5-16E3 | Biolegend |
| TGF-β | TW7-16B4 | Biolegend |
| FOXP3 | MF-14 | Biolegend |
| Granzyme b | QA16A02 | Biolegend |
| Perforin | S16009A | Biolegend |
| Tbet | 4B10 | Biolegend |
| CD28 | 37.51 | Biolegend |
| CD57 | QA17A04 | Biolegend |
| KLRG1 | 2F1/KLRG1 | Biolegend |
| βgalactosidase | 40-1a | Santacruz |

**Table 2:**
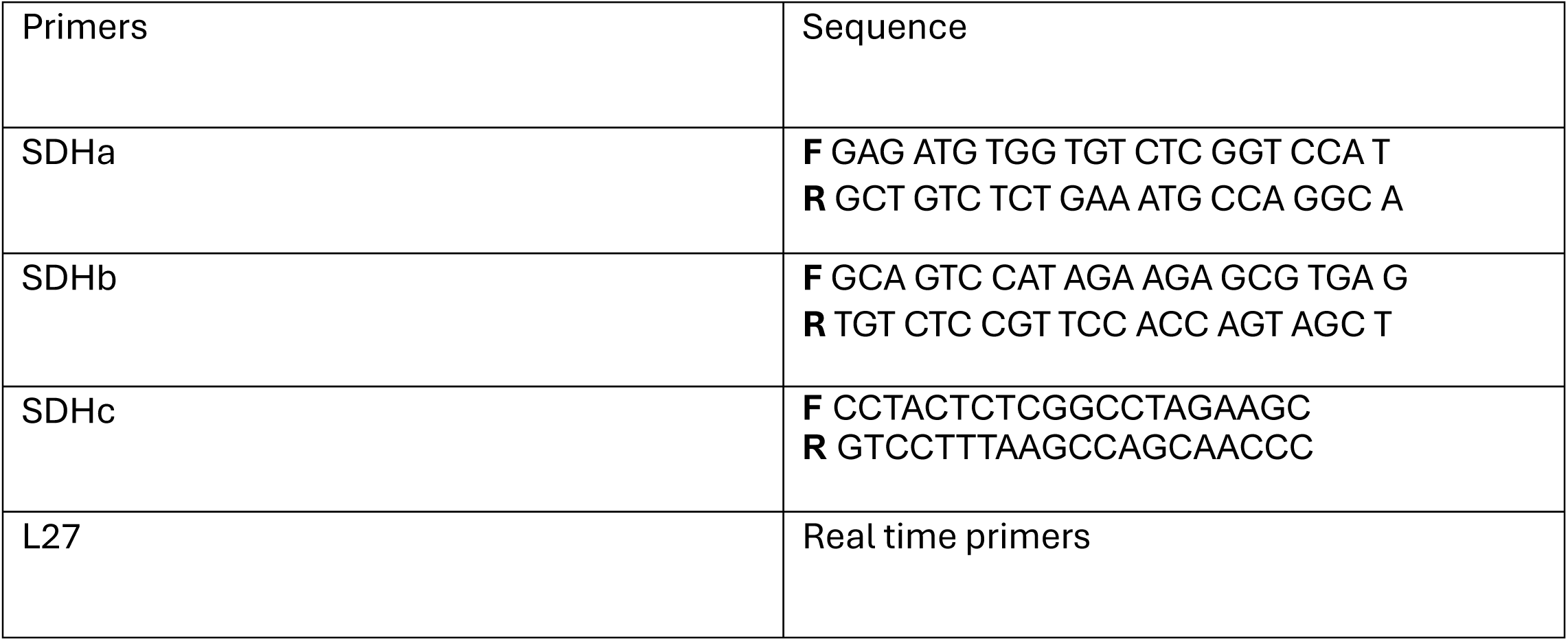
List of primers.

## Notes

### Competing Interest Statement

The authors have declared no competing interest.

