## Supplementary figure legends and Figures for "Aging-Driven Immunosuppression: The Role of Tregs in the Ovarian Tumor Microenvironment"

Supplementary Figures:

**Figure S1: Aging exacerbates preclinical EOC models:** ID8-luc2 cells were injected in young and old EOC mice, (A). Representative BLI images at 4 weeks of tumor inoculation, (B) Bar graph of BLI quantification of images in A. (C) Average weekly body weight progression in ID8-luc2 mice. (D) Average weekly body weight progression in $ID8^{p53-/-}$ mice. ∗∗∗$p < 0.001$, OT compared with YT group by Student's *t* test.

**Figure S2: Aging enhances tumor promoting metabolic growth factors and Senescence Associated Secretory Phenotype associated inflammation:** (A–X) Measurement of (A, E) Insulin, (B, F) IGF-1, (C, G) Adiponectin, (D, H) Leptin, (I, M) IL-4, (J,N) MCP-1, (K,O) IL-10, (L, P) GMCSF, (Q, U) IL-1β, (R,V) TNFα, (S, W) IL-6, (T, X) IFN γ, and in ascites collected at week 5 from $ID8^{p53-/-}$ and $ID8^{p53-/-, BRCA 1-/-}$ EOC bearing mice by ELISA (n = 3).∗$p < 0.05$, ∗∗$p < 0.01$, ∗∗∗$p < 0.001$, OT compared with YT group by Student's *t* test.

**Figure S3: Aging induces differential systemic T cell response in response to EOC:** $ID8^{p53-/-}$ EOC injected in Y.C, Y.T or O.C, O.T (n = 10/group), (A–I) (A) A representative t-SNE visualization of markers after gating on single, live, $CD45^+$ $CD3^+$, $CD4^+$, $CD8^+$, IFNγ, Granzyme B (Grz B), perforin and FOXP3 expression. (B) Heatmap represents marker expression of the main T cell subsets in individual samples. Immune profiling was performed in blood of 5 individual mice per group. Violin plots represent the percentage of T cell subsets (C) CD4+, (D) CD4+IFNγ+, (E) CD4+ CD25+ FOXP3 (F) CD8+, (G) CD8+IFNγ+, (H) CD8+Grzb+, (I) CD8+Perforin+, (J) CD11b+ F4/80, (K) F4/80+CD38, (L) F4/80+ EGR2, (M) F4/80+ CD206, (N) CD38/EGR2, (O) CD38/CD206, (P) CD38+iNOS, (Q) EGR2+Arg1, (R) CD206+ Arg1, (S) CD11b+ GR1. The experiment was repeated twice in two different sets of mouse experiments. ∗$p < 0.05$, ∗∗$p < 0.01$, ∗∗∗$p < 0.001$.

**Figure S4: Aging diminishes the T cell response in EOC TME:** ID8$^{p53-/-}$ EOC injected in Y.T or O.T (n = 10/group), Immune profiling was performed in ascites of 5 individual mice per group (A–R) (A) A representative t-SNE visualization of markers after gating on single, live, CD45$^+$ CD3$^+$, CD4$^+$, CD8$^+$, IFNγ, Granzyme B, perforin and PD1 expression. (B) Heatmap represents marker expression of the main T cell subsets in individual samples. Violin plots represent the percentage of T cell subsets (C) CD8$^+$, (D) CD8+IFNγ+, (E) CD8+GrzB+, (F) CD8+perforin, (G) CD8+ PD1. (H) A representative t-SNE visualization of markers after gating on single, live, CD45$^+$ CD3$^+$, CD4$^+$, IFNγ, and Tbet expression. (I) Heatmap represents marker expression of the main T cell subsets in individual samples. Violin plots represent (J) CD4, (K) CD4$^+$ IFNγ+ (L) CD4+Tbet. (M) A representative t-SNE visualization of markers after gating on single, live, CD45$^+$ CD3$^+$, CD4$^+$, CD25+, FOXP3, IL-10, TGF-β and PD1 expression. (N) Heatmap represents marker expression of the main T cell subsets in individual samples. Violin plots represent the percentage of T cell subsets (O) FOXP3, (P) FOXP3+ TGF-β, (Q) FOXP3+ IL-10, (R) FOXP3+PD1 in the tumors. The experiment was repeated twice in two different sets of mouse experiment. ∗$p < 0.05$, ∗∗$p < 0.01$, ∗∗∗$p < 0.001$, OT compared to YT group by Student's *t* test.

**Figure S5: Aging diminishes the intra-tumor T cell response in EOC:** ID8$^{p53-/-, BRCA\ 1-/-}$ EOC injected in Y.T or O.T (n = 10/group), Immune profiling was performed in tumors of 5 individual mice per group (A–R) (A) A representative t-SNE visualization of markers after gating on single, live, CD45$^+$ CD3$^+$, CD4$^+$, CD8$^+$, IFNγ, Granzyme B (Grz B), perforin, Tbet and FOXP3. (B) Violin plots represent the percentage of T cell subsets (B) CD4+ (C) CD4+IFNγ, (D) CD4+ Tbet, (E) CD25+ FOXP3, (F) FOXP3+IL-10, (G) FOXP3+ TGFβ, (H) CD8+, (I) CD8+ IFNγ, (J)

CD8+ Granzyme b, (K) CD8+ Perforin. The experiment was repeated twice in two different sets of mouse experiments. $**p < 0.01$, $***p < 0.001$, OT compared to YT group by Student's *t* test.

**Figure S6: Tregs are key in promoting EOC:** (A) Experimental plan showing the Treg depletion strategy. (B)Kaplan–Meier graph indicating overall survival (n = 10), p = 0.002 by Gehan-Breslow-Wilcoxon test. (C) Average ascites accumulated in O.T, O. Treg dep. Immune profiling was performed in blood (D) A representative t-SNE visualization of markers after gating on single, live, CD45, CD4, CD8, IFNγ, Granzyme B (Grz B), perforin and PD1. Violin plots represent the percentage of senescence markers on T cell subsets (E) CD4+ (F) $CD4^+$ IFNγ+, (G) FOXP3, (H) FOXP3+ IL-10, (I) FOXP3+ TGFβ, (J) Heatmap represents marker expression of the main T cell subsets in individual samples. (K) $CD8^+$, (L) CD8+IFNγ+, (M) CD8+perforin, (N) CD8+GrzB+, (O) CD8+ PD1, (P) CD8+ KLRG1, (Q) CD8+ βGal, (R) CD4+ CD57, (S) CD4+CD28 (T) CD4+ KLRG1, (U) CD4+βGal, (V) CD4+CD57, (W) CD4+CD28. Immune profiling was performed in 4 individual mice per group. $*p < 0.05$, $**p < 0.01$, $***p < 0.001$, by one-way ANOVA, followed by Sidak multiple comparison test.

**Figure S7: Tregs from aged EOC mice prefer OXPHOS for immunosuppression:** $ID8^{p53-/-, BRCA\ 1-/-}$ EOC cells were injected in Y.T or O.T , FOXP3+ cells were isolated from the spleens of EOC young and old tumor at 5 weeks and subjected to Seahorse analysis and energy targeted metabolomics. (A) Oxygen consumption rate (OCR) was assessed in real-time using an XFe Seahorse analyzer as described in methods. Port injections of (1) oligomycin, (2) FCCP, and a combination of (3) rotenone-antimycin were given. (B) The bar graph represents basal and stressed OCR (n = 3). (C) Extracellular acidification rate (ECAR) was measured with port injections of (1) glucose, (2) oligomycin, and (3) 2-DG. (D) The bar graph represents basal and stressed ECAR (n = 3). $***p < 0.001$, OT compared to YT group by Student's *t* test.

**Figure S8: Altered Glycolytic Metabolite Profiles in Young and Aged Regulatory T Cells:** Tregs were isolated from control and tumor bearing young and old mice and were processed for Glycolysis metabolites. (A-G) Targeted analysis of the major glycolysis metabolites was performed to assess the levels of various metabolites in pooled xenografts ($n = 3$) in triplicates.

**Figure S9 Tumor derived succinate promotes Treg function in aged EOC mice:** Tregs isolated from young EOC mice and cultured with and without succinate (Succinate 100 µm,1 mm,5 mm). Bar graph represents the percentage of A. CD25+FOXP3 cells. B. FOXP3+IL10 and C. FOXP3+TGFβ. D. CD4+CFSE from Tregs cocultured with naïve CD4 T cells in presence and absence of succinate (100 µm,1 mm,5 mm). Tregs from young EOC mice were exposed to young and old patient serum and flow analysis was performed Bar graph represents (E) %CD25+ FOXP3, (F) FOXP3+ 1L-10, (G) FOXP3+ TGF-β. Tumors cells from aged EOC mice were treated with CPI-613 and the tumor condition media (TCM) and CPI-613+TCM media was exposed to Tregs from young EOC mice. Bar graph represents the percentage of (H) succinate levels in the tumor cells after treatment with CPI-613. (I) CD25+FOXP3, (J) FOXP3+ IL-10, (K) FOXP3+ TGFβ. (L) CFSE proliferation histograms and bar graph representing CD4 proliferation in the presence of Tregs exposed to TCM and CPI-613+TCM. Flow analysis was performed in the co-culture experiment, and bar graph represents senescence markers (M) CD4+ CD28, (N) CD4+ KLRG1, (O) CD4+ CD57, (P) CD4+ β-Gal. Bioenergetics using seahorse was performed on Tregs exposed to TCM and CPI-613+TCM (Q) Oxygen consumption rate (OCR) was assessed in real-time using an XFe Seahorse analyzer as described in methods. Port injections of (1) oligomycin, (2) FCCP, and a combination of (3) rotenone-antimycin were given. (R) The bar graph represents basal and stressed OCR (n = 3). (S) Extracellular acidification rate (ECAR) was measured with port injections of (1) glucose, (2) oligomycin, and (3) 2-DG. (T) The bar graph represents basal and stressed ECAR (n = 3). ***$p<0.001$, **$p<0.01$, *$p<0.5$.

**Figure S10: Succinate inhibition reduces Treg function:** Tumors cells from aged EOC mice were treated with AA6 and the tumor condition media (TCM) and TCM+AA6 media was exposed to Tregs from young EOC mice. Bar graph represents the percentage of (A, E) CD25+ FOXP3, (C, H) FOXP3+ IL-10, (B, F) FOXP3+ TGFβ and (D, G) PD1.***$p<0.001$, TCM+AA6 compared with TCM group by Student's *t* test.

### Supplementary figures

**Figure S1: Aging exacerbates preclinical EOC models**

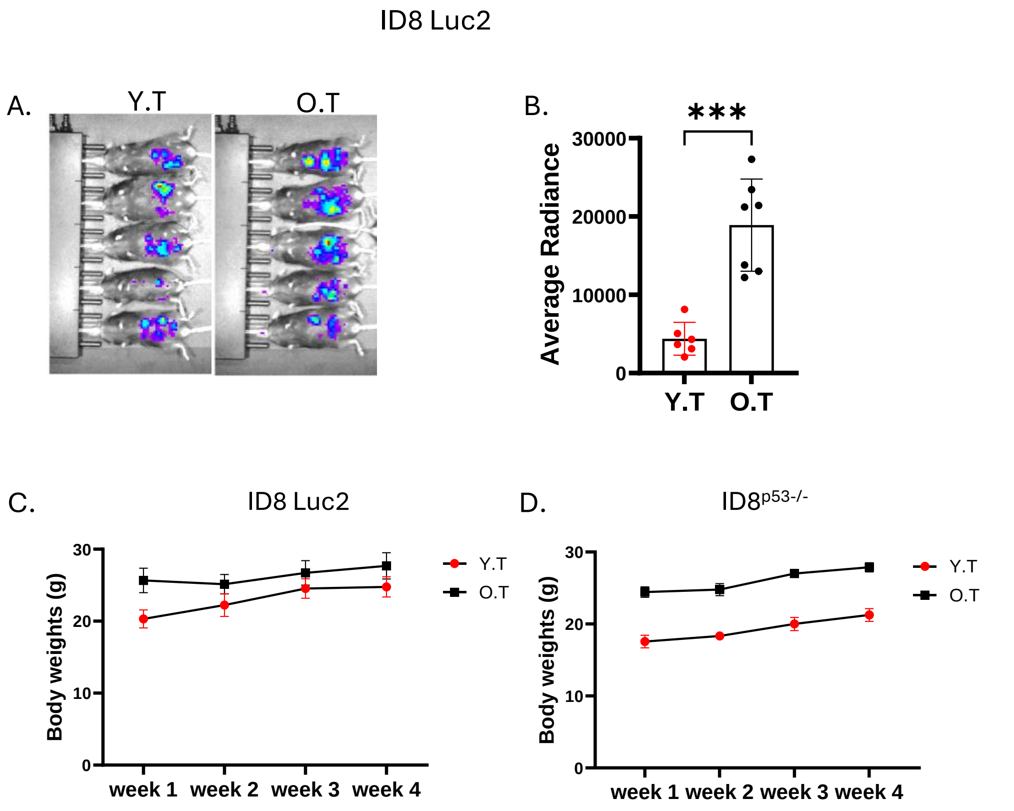

**Figure S2: Aging enhances Senescence Associated Secretory Phenotype in EOC.**

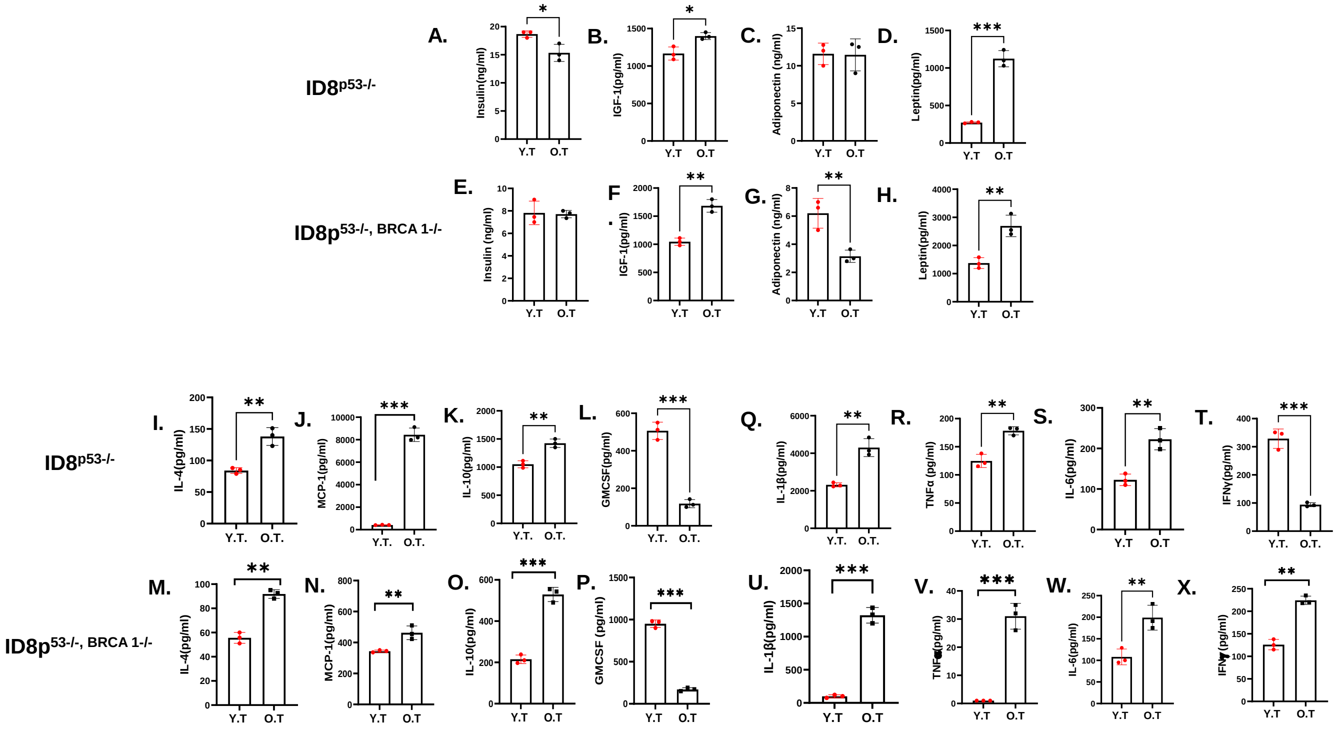

### S3: Aging induces differential systemic immune responses in response to EOC

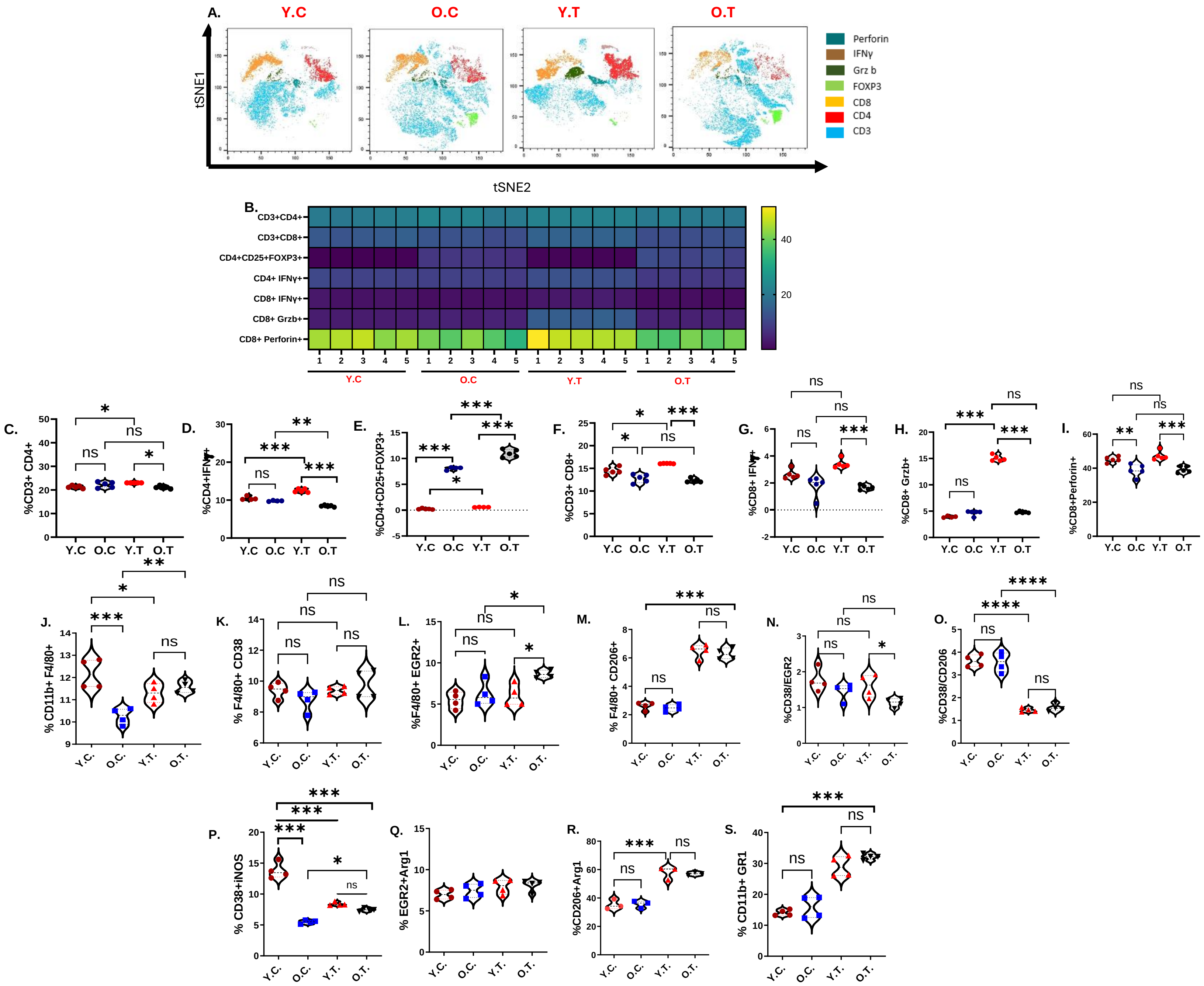

### Figure S 4: Aging induces Differential T cell response in Ovarian TME.

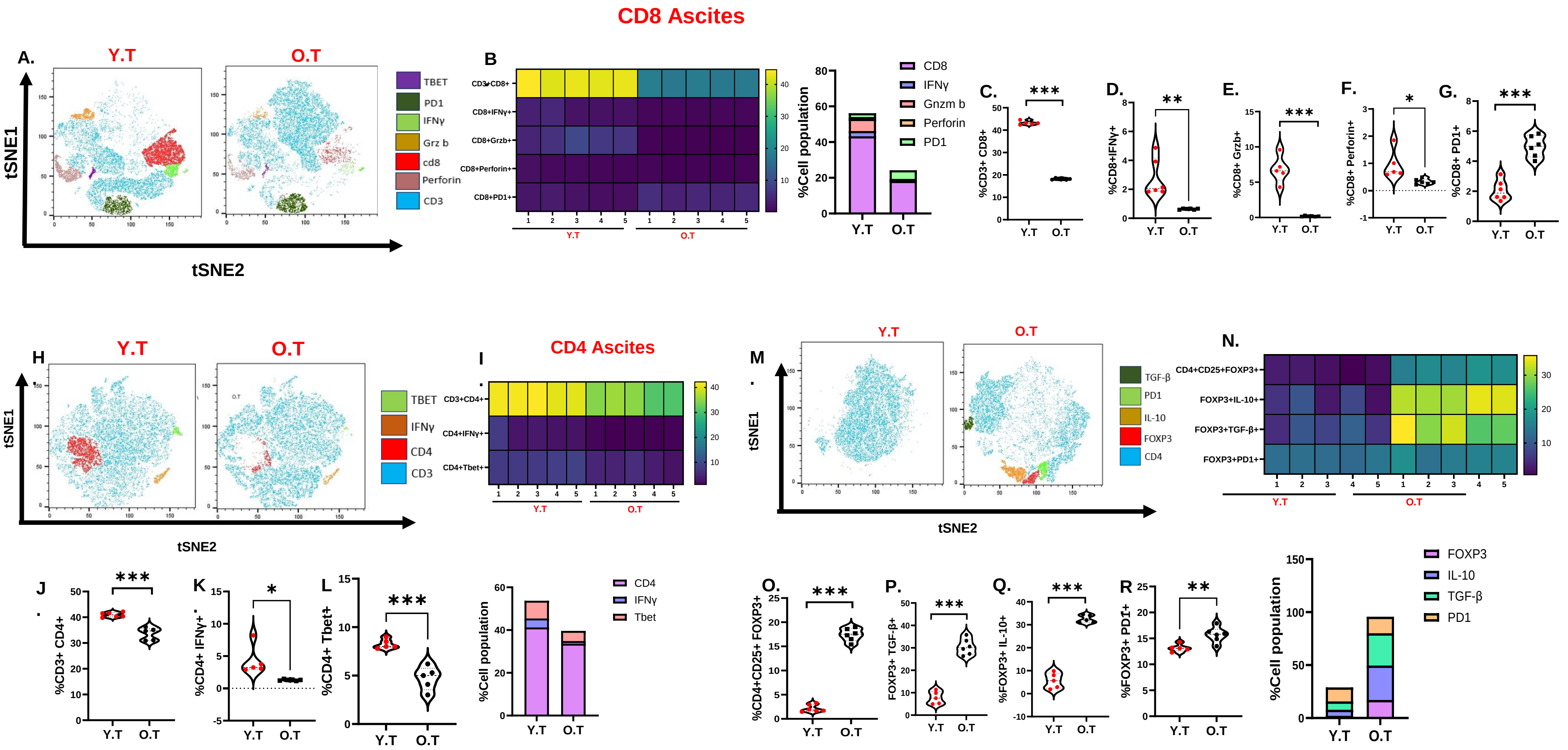

#### Figure S5: Aging diminishes the intratumor T cell response in ID8 BRCA1-/- injected EOC mice.

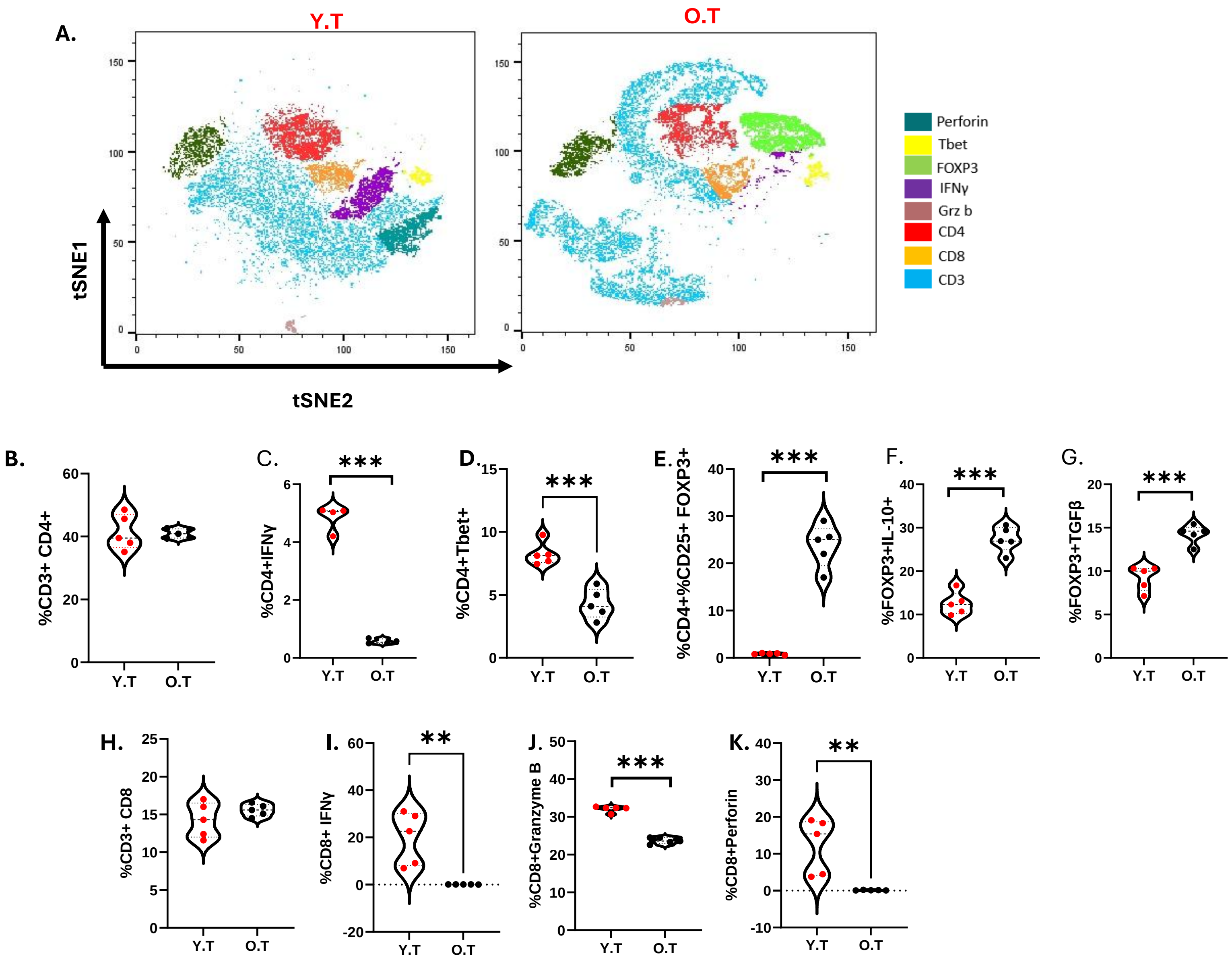

### Figure S6: Tregs are key in promoting EOC

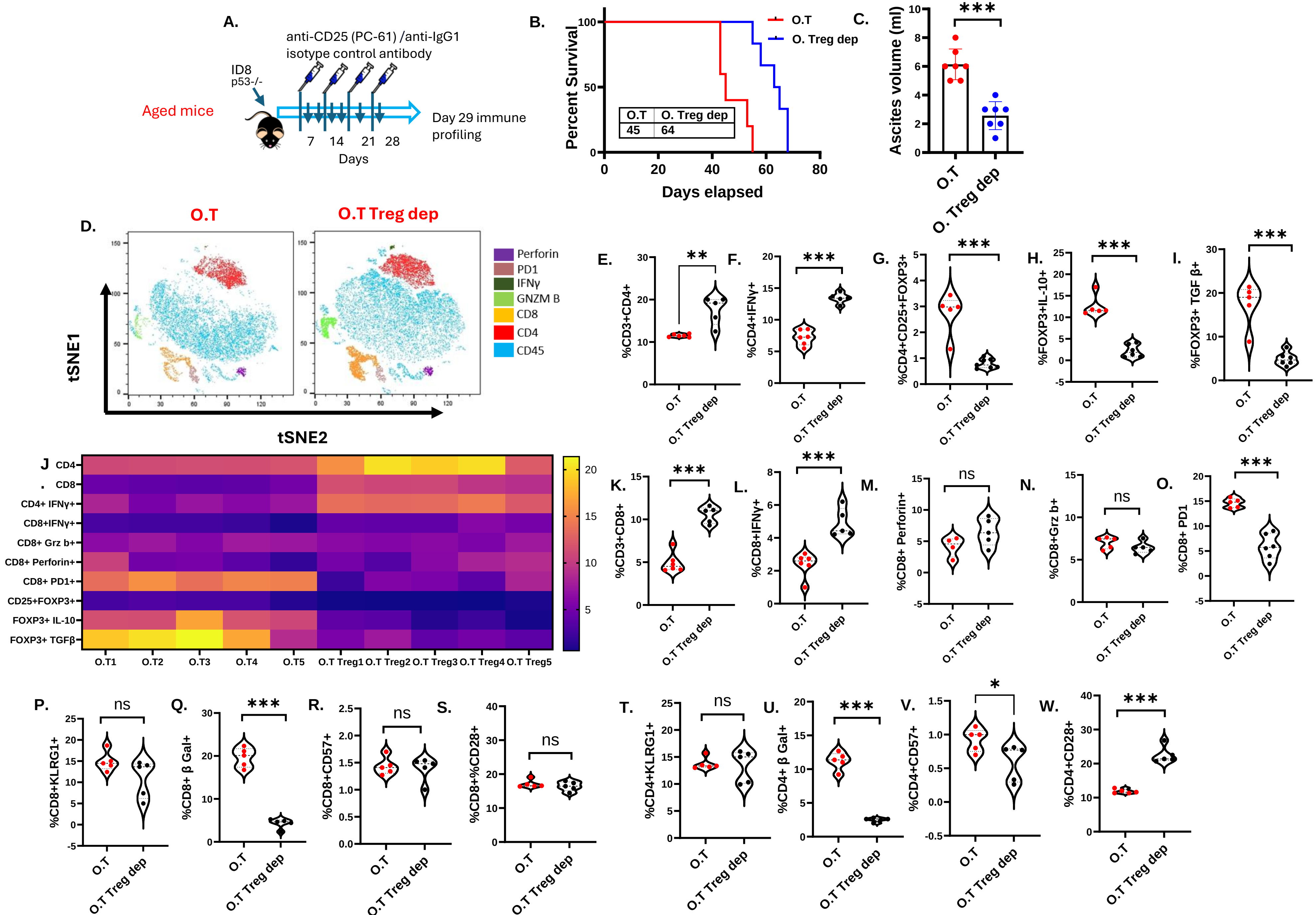

#### Fig S7: Tregs from aged EOC mice prefer OXPHOS for immunosuppression.

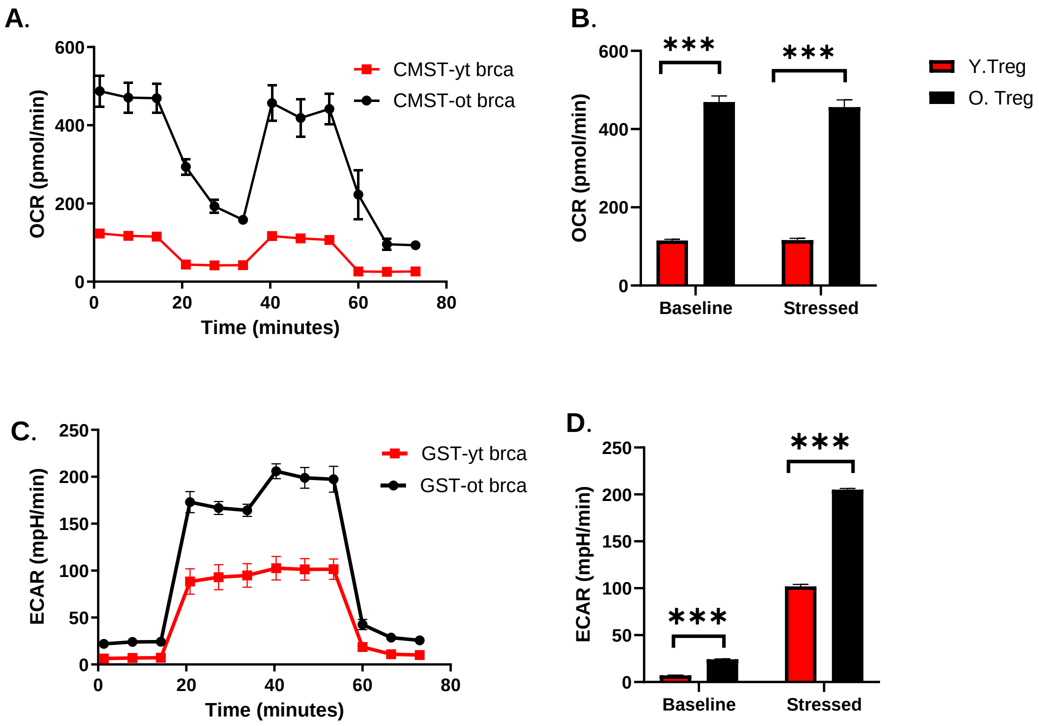

#### Fig S8: Glycolysis metabolites in Tregs of young and old mice

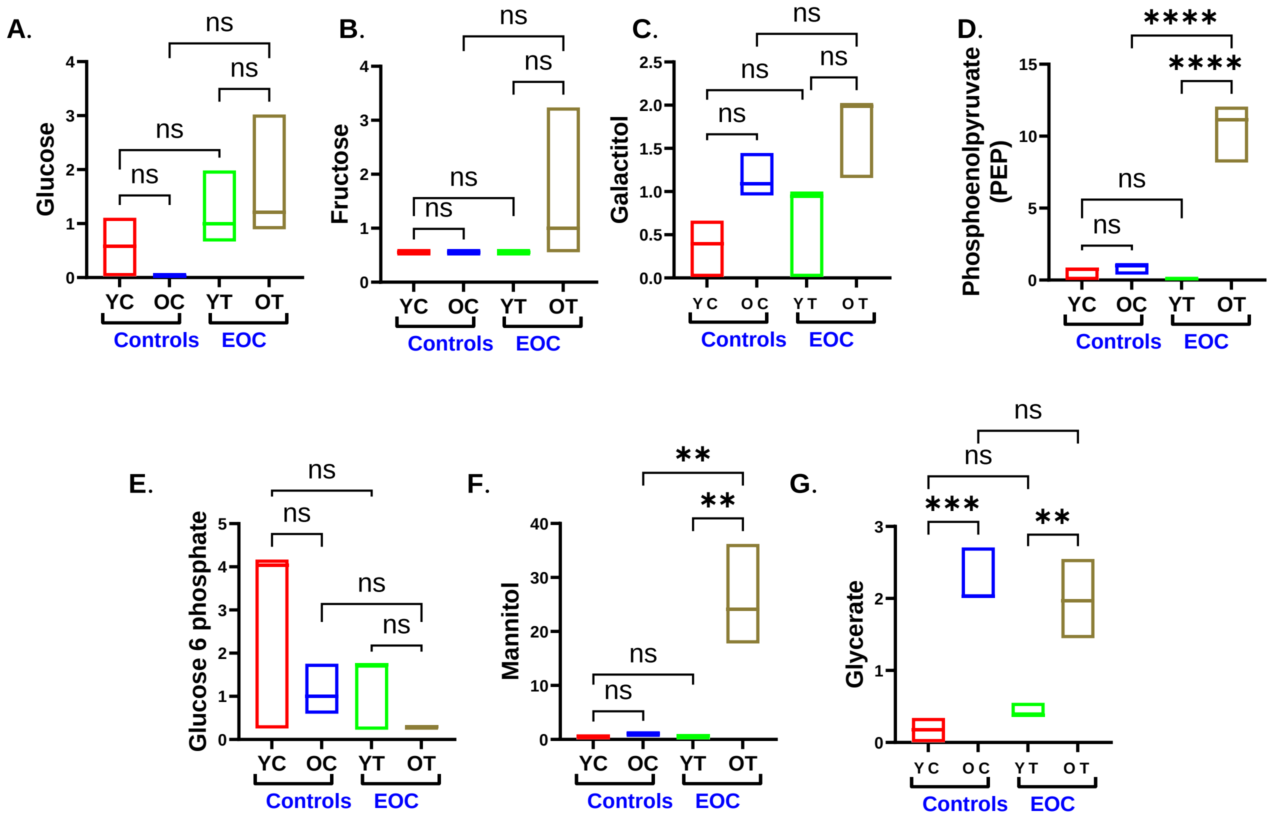

### Fig S9: Tumor derived succinate promotes Treg function in aged EOC mice

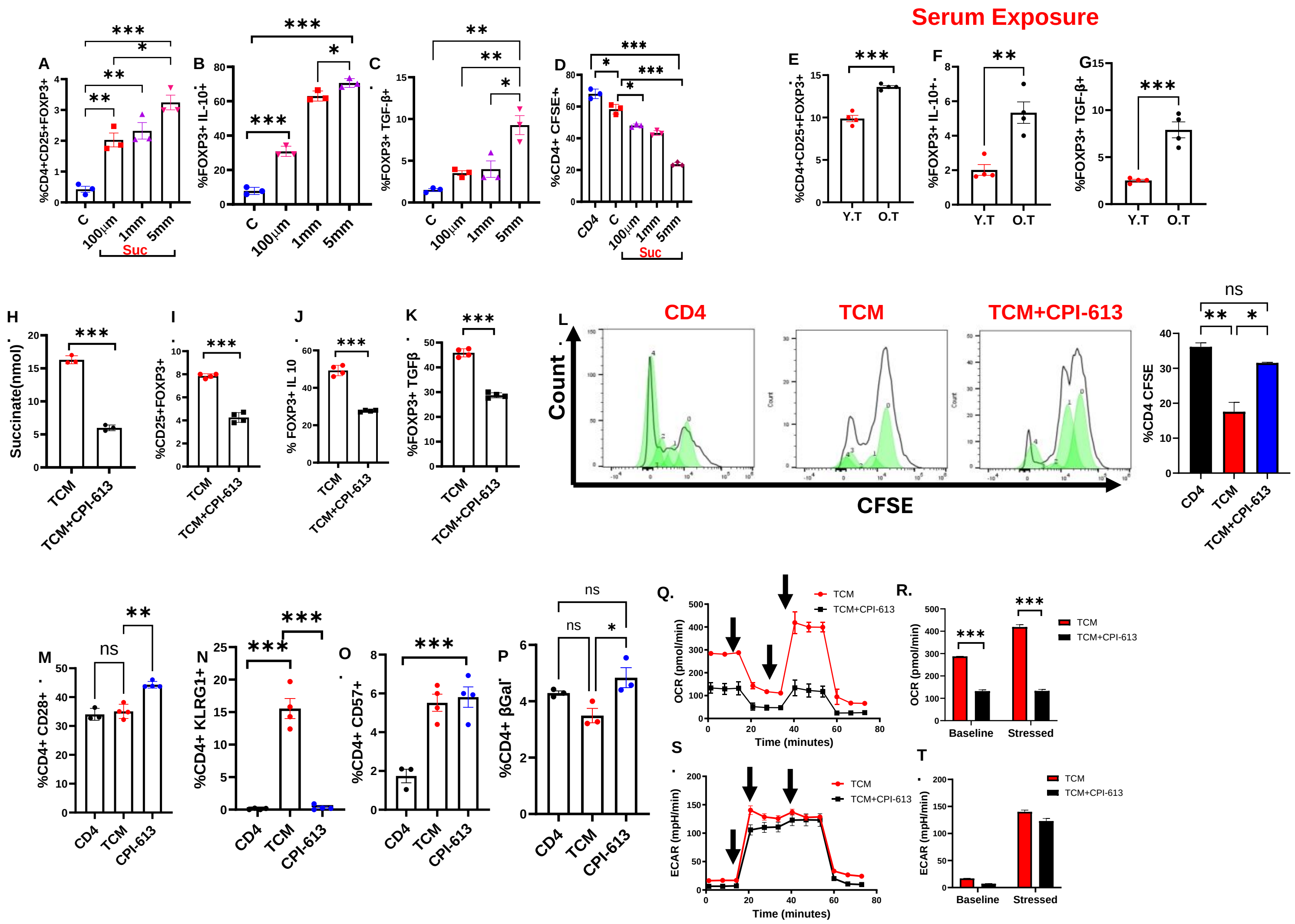

**Fig S10: Inhibition of succinate synthesis decreased Treg function**

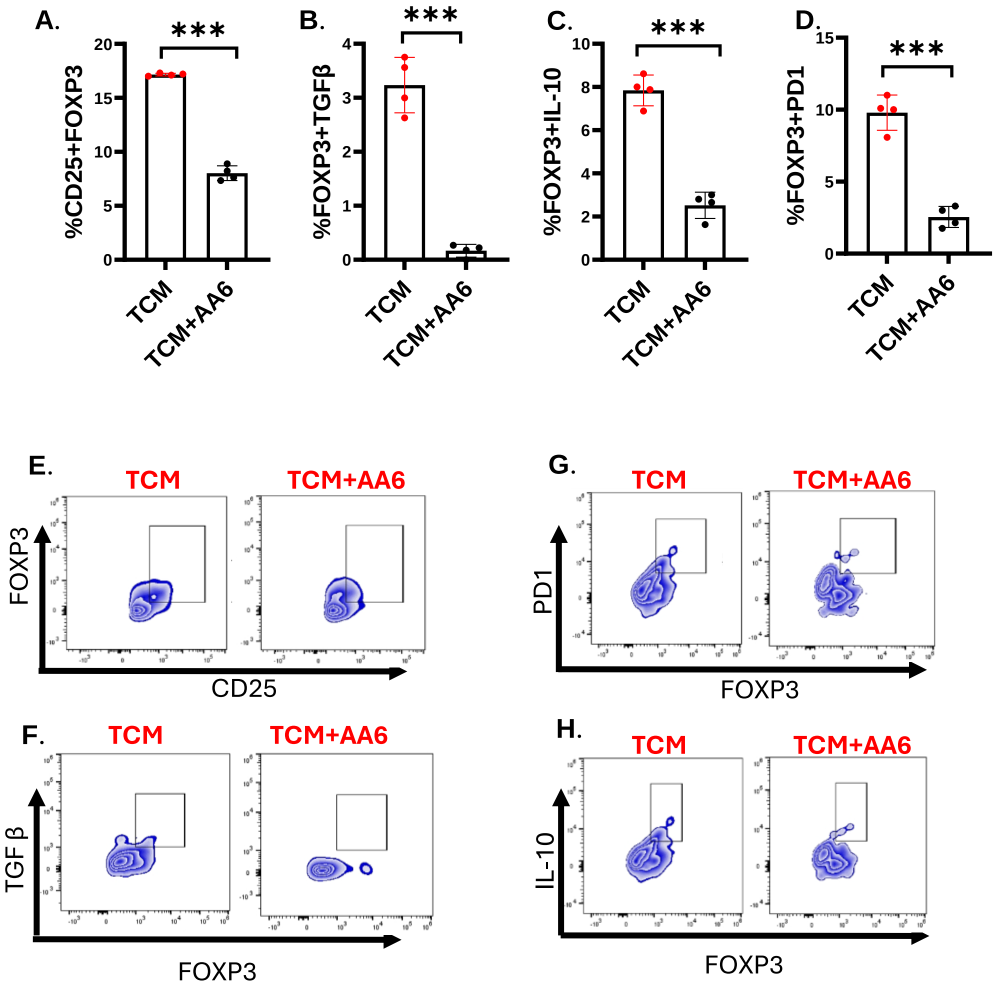
